# Identifying neuropeptides in *Hydra*: a custom pipeline reveals a non-amidated regulator of muscle contraction and other new members

**DOI:** 10.1101/2025.03.12.642944

**Authors:** Pranav Prabhu, Puli Chandramouli Reddy

## Abstract

Neuropeptides play a critical role in neurotransmission and organismal development. Members of phylum Cnidaria, with a diffused nervous system, are one of the earliest divergent animals and might provide insights into the fundamentals of the emergence of neuronal communications. The neuropeptide diversity in *Hydra* (a cnidarian model) has been extensively studied using various strategies, each with certain limitations. Here, we have developed an *in silico* pipeline which identified both reported peptides and many new potential candidates. A comparative analysis within Cnidaria suggests a rapid divergence of neuropeptides which might be involved in complex behaviors. We identified new *Hydra* neuropeptides that belong to the RFamide and PRGamide families and a novel class of peptides lacking amidation (LW-peptides). A detailed expression and functional analysis of a new LW-peptide indicates its role in the longitudinal contraction of *Hydra* polyps. This study provides compelling evidence for the existence of intricate peptidergic communication in early neuronal circuits. The extensive diversity of neuropeptides within this phylum underscores their rapid evolutionary adaptability. This current pipeline also proves to be simple and adaptable to perform neuropeptide identification in other multicellular organisms.

## Introduction

Peptidergic signaling is one of the oldest methods of communication employed by early nervous systems. Neuropeptides are short amino acid chains typically encoded in multiple units in a larger precursor. Proteins harboring neuropeptides are post-translationally cleaved by endopeptidases following the removal of the protein’s signal sequence. These fragments undergo further modifications yielding mature peptides released from secretory vesicles into the extracellular space. Common modifications include C-terminal amidation and N-terminal pyroglutamate conversion that prevent enzymatic degradation [1].

Cnidaria—a diploblastic sister group to Bilateria—is a large phylum of early diverging aquatic animals characterized by their stinging nematocytes. They possess diffuse nerve nets primarily peptidergic in nature [2,3] that control various physiological processes including muscular movement, feeding behavior, sensory activity, and reproduction [4–7]. Studying cnidarian neuropeptides can provide insights into the evolution of nervous systems and their roles in controlling basal organismal behaviors. The bioactive nature of such peptides and their conservation across Eumetazoa also make them intriguing molecules for pharmacological applications.

There has been significant inquiry into *Hydra* neuropeptides due to its key phylogenetic position at the base of Metazoa. Over the decades, 5 major families have been identified: the RFamides, KVamides, FRamides, GLWamides, and PRXamides [8], in addition to peptides such as NDA-1 [9]. While RFamides and GLWamides are conserved in other metazoans [10], some such as KVamides are species-specific [11]. *Hydra* neuropeptides were initially described through peptidomics and other experimental means [12]. Recent studies have attempted to computationally predict cnidarian neuropeptides using sequence properties and common motifs [13–19]. In this paper, we present an improved neuropeptide prediction pipeline that incorporates recent single-cell RNA sequencing data and modified parameters to identify new neuropeptide candidates in three model cnidarians: *Hydra vulgaris, Nematostella vectensis*, and *Clytia hemisphaerica*. We characterize the expression of three new *Hydra* neuropeptide precursors, one of which encodes a unique non-amidated neuropeptide that induces body column and tentacle contraction in *Hydra*.

## Results

### A custom script identifies new neuropeptide precursors

To comprehensively identify neuropeptide precursors, we used predicted protein sequences from *H. vulgaris* AEP, *C. hemisphaerica*, and *N. vectensis* [20–22] containing signal peptides, lacking transmembrane regions, and which did not encode functional domains present in the Pfam database. These were scored using our custom script to extract sequences with known C-terminal cleavage sites (KK, RK, KR, RR, GK, GR) and N-terminal motifs (Q, XP) within a specified range that could potentially encode peptides. Sequences with non-neuropeptide homologs were removed. We then filtered and clustered sequences expressed in neuronal populations using single-cell gene expression data to obtain a manually curated list of putative precursors (Fig 1A). Combined similarity clustering resulted in three large clusters: RFamides and GLWamides, the two conserved peptide families and their derivatives, and a cluster of random assorted peptides (Fig 1B, S2 Fig). Interestingly, another large cnidarian neuropeptide group of PRXamides present in all three organisms did not cluster together.

**Figure 1.**
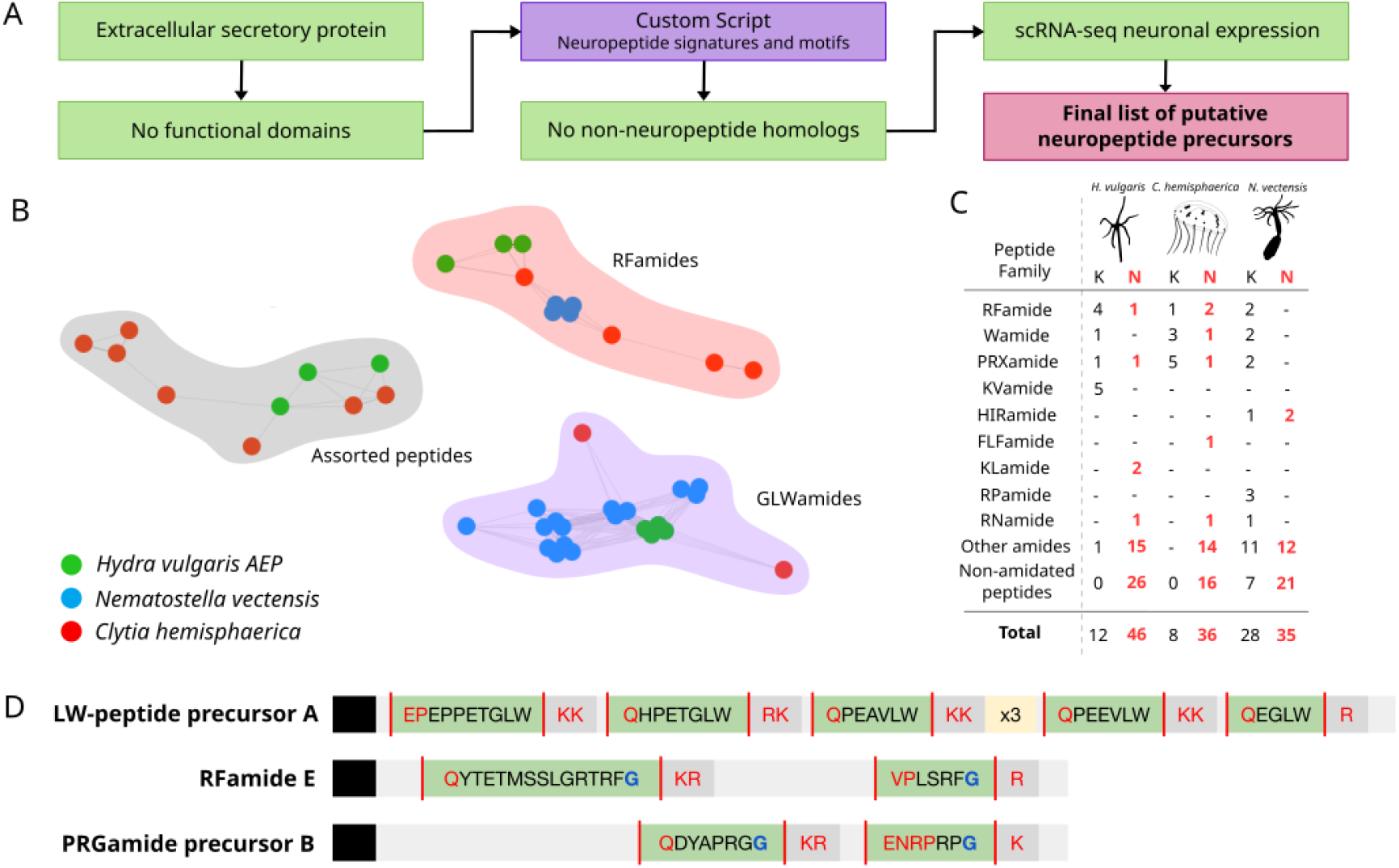
A custom prediction script identifies new cnidarian neuropeptide precursors. (A) A simplified flowchart of the pipeline developed to identify neuropeptide candidates. (B) Network map of selected recovered clusters from combined similarity clustering by CLANS. (C) Table summarizing the number of newly identified neuropeptide proteins ‘N’ in black font alongside their known counterparts ‘K’ in red font for all three organisms. (D) Sequence structure of the three *Hydra* neuropeptide precursors characterized in this study.

In addition to neuropeptides reported in previous studies, we identified many new precursors in all three cnidarians (Fig 1C, S1 Table, S1 Data). In *Clytia*, genes encoding various PRXamides, RFamides, KFamides and GLWamides are known [16,23]. Here, report another precursor encoding different GLWamides and an RWamide. Additionally, genes encoding a new family of FLFamides, as well as FAFamides, RYamides, RIamides, FAVamides, and ANamides, were also identified (Fig 1C). We also found genes encoding peptide repeats with common C-terminal motifs, including QQ-peptides, KIM-peptides, MLS-peptides, and LT-peptides. A comprehensive list of *Nematostella* peptides has already been assembled from many previous neuropeptide studies [17–19]. Adding to this list, we identified a new precursor containing a variety of neuropeptides, including EFamide, LMamide, and NAamide (Fig 1C). A set of sequences also encoded previously unreported GLWC and GFWC-peptides that show significant similarity with other GLWamide and WGC-peptide precursors (Fig 1B). In total, we were able to predict 36 new neuropeptide genes in *Clytia* and 35 genes in *Nematostella*.

We identified 46 new *Hydra* genes encoding potential neuropeptides, more than both other cnidarians. We chose three most suitable candidates to characterize further: G016204, G004106, and G009409 (Fig 1D, S3 Fig). The first, which we term Preprohormone E, encodes two putative RFamides and is present at a different locus from the known RFamide preprohormones, but on the same chromosome. G004106 (referred to as PRGamide precursor B) contains two reported but uncharacterized PRXamides. G009409, which we named LW-peptide A, encodes multiple LW-peptides, all of which lack the typical amidation residue. Other identified precursors encoded putative peptides such as INamide, KLamide, KIamide, and CGH/Y-peptides, among many others (Fig 1C).

### Gene expression analyses of putative *Hydra* neuropeptide genes

We first performed *in situ* hybridization to confirm whether these genes are neuropeptide precursors. All three exhibit clear neuronal expression and have differing expression patterns corresponding to particular subpopulations. LW-peptide precursor A is expressed in neurons across the polyp with higher intensity in the head and foot regions, including the basal disk. However, it appears to be completely absent in the tentacles, with expression restricted to a clear ring around each tentacle at its base (Fig 2A-D). PRGamide precursor B exhibits expression in neurons across the body column but is absent in the tentacles and below the peduncle. There is also a clear ring of expression around the mouth. Many of the body neurons appear to be bipolar, indicating these are likely ganglion cells, but there are others spanning the ectoderm-endoderm boundary that are likely sensory neurons [24] (Fig 2E-I). RFamide preprohormone E is primarily expressed in tentacular neurons, in addition to a few peduncular neurons (Fig 2J-M).

**Figure 2.**
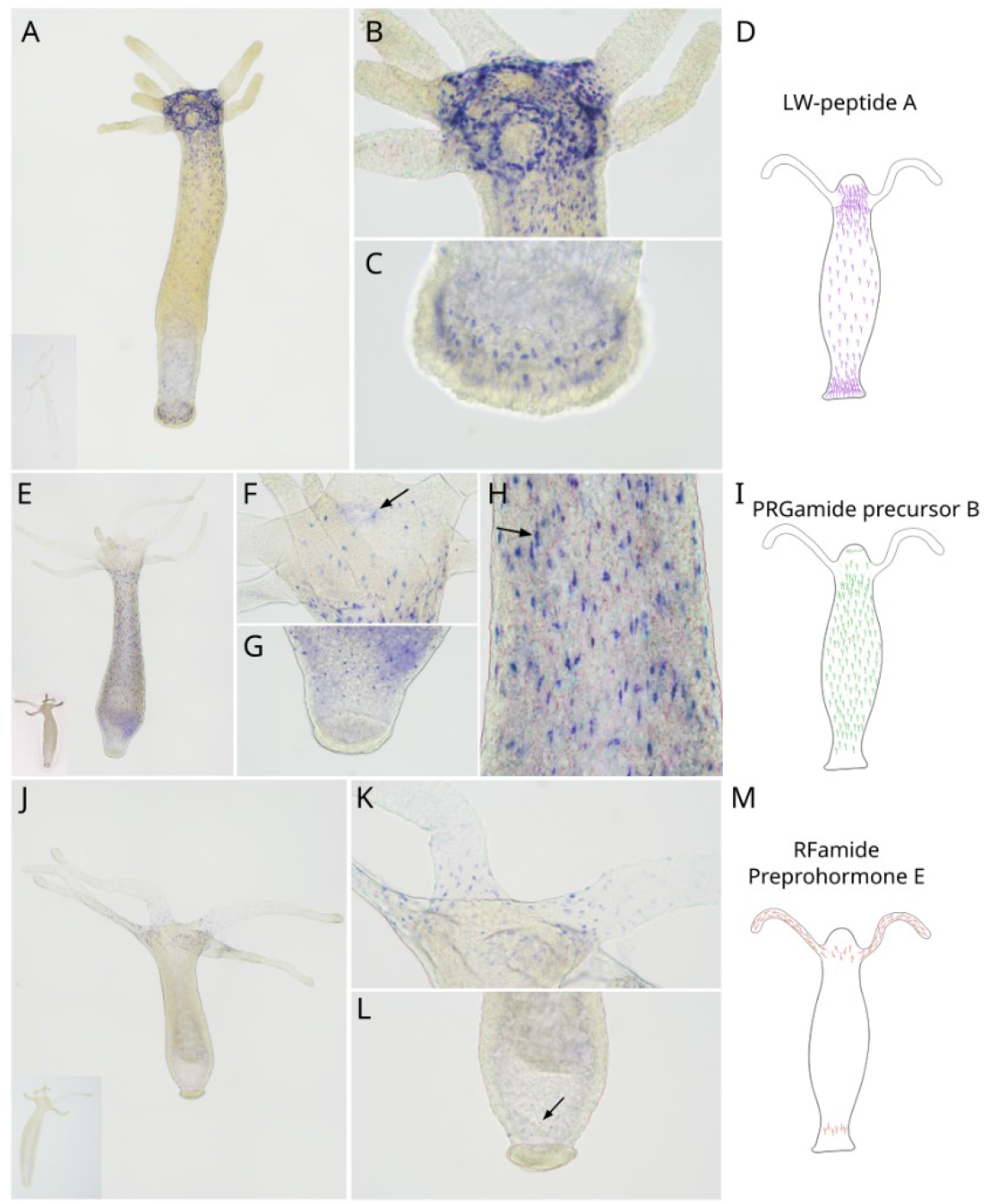
Gene expression analyses of putative *Hydra* neuropeptide genes using whole mount *in situ* hybridization. (A-C) WISH images displaying the expression of LW-peptide precursor A across (A) the whole polyp, (B) head region, and (C) foot region. (E-H) WISH images displaying the expression of PRGamide precursor B across (E) the whole polyp, (F) head region with the arrow pointing to the mouth, (G) foot region, and (H) the gastric region with the arrow pointing to a bipolar neuron. (J-L) WISH images displaying the expression of RFamide Preprohormone E across (J) the whole polyp, (K) the hypostome and tentacles, and (L) the foot region with the arrow pointing to the few cells exhibiting RFamide Preprohormone E expression. Inset images in A, E and J are sense controls. Images (A, E, J) were captured at 100x magnification and stitched while the others were captured at 100x and stacked. (D, I, M) Schematic representations of each gene’s expression pattern across the polyp.

### A non-amidated LW-peptide induces contraction in *Hydra*

Unlike known *Hydra* neuropeptides, none of the putative peptides encoded by LW-peptide precursor A contain the C-terminal Glycine residue required for amidation. Experiments using cnidarian GLWamides have demonstrated that this amidation is required for their bioactivity [25–27]. We were interested in whether LW peptides could induce any specific behavior on exogenous addition and whether this was similar to GLWamides. On treating glycocalyx-removed polyps using 0.1mM of LWa1 peptide (amino acid sequence: EPEPPETGLW), we observed immediate body column contraction followed by tentacular contraction compared to the control. This effect was prolonged, with many polyps remaining contracted for an hour after treatment. Within just 10 seconds, there was a significant percentage reduction in body length and average tentacle length per polyp (Fig 3). This clearly indicates that LWa1, a non-amidated peptide, specifically induces contraction in *Hydra*, making this the first record, to our knowledge, of a bioactive non-amidated neuropeptide in Cnidaria and the first such putative LW-peptide in Metazoa. Additionally, unlike other *Hydra* GLWamides that are known to induce bud detachment, this peptide showed no propensity for the same (S5 Fig).

**Figure 3.**
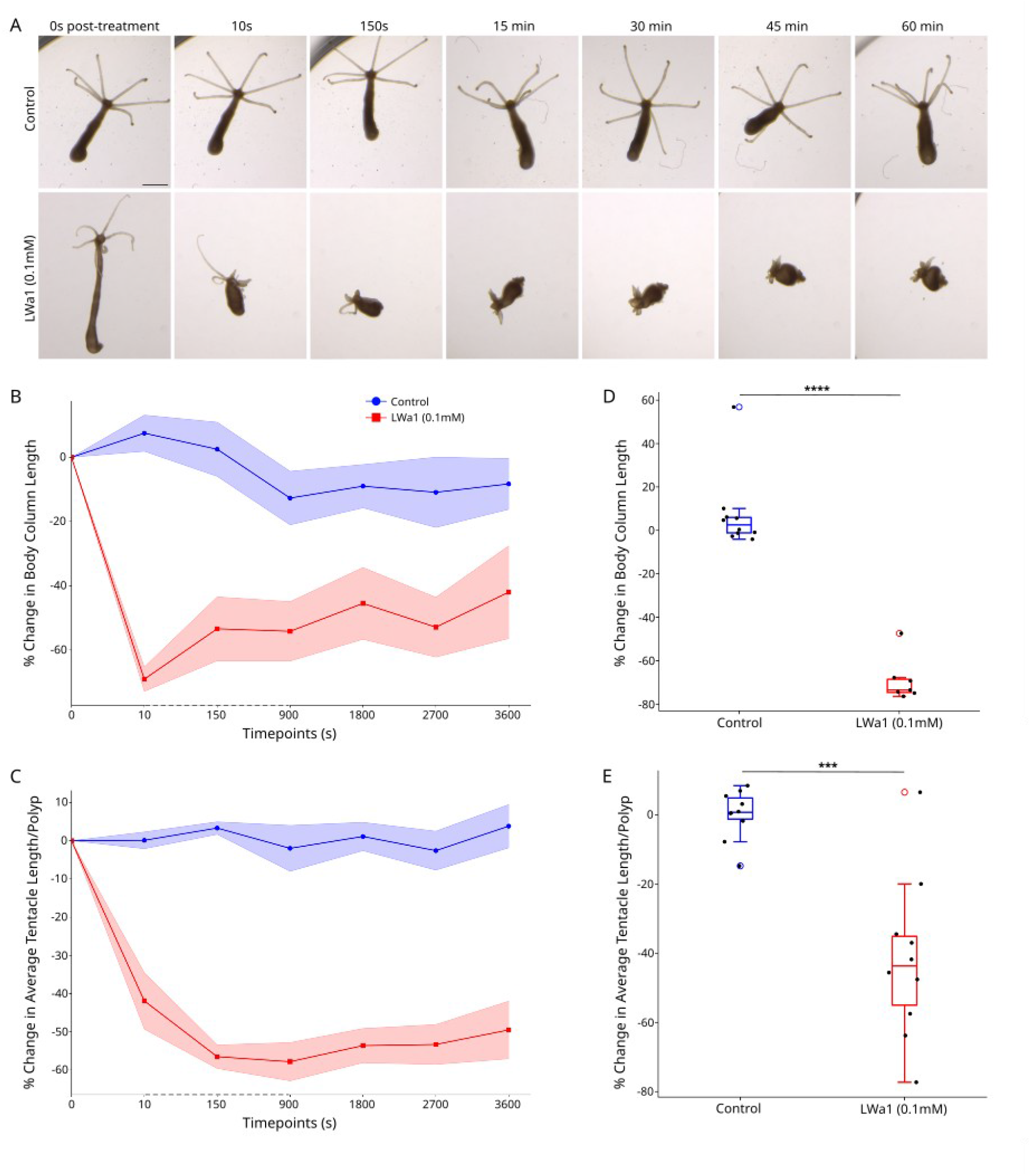
LWa1 peptide induces muscle contraction in *Hydra*. (A) Snapshots of polyp behavior at various timepoints post-treatment with either the control or 0.1mM Lwa1. (B-C) Trendlines of percentage change in (B) body column length (n=7-10) and (C) average tentacle length per polyp (n=min.7) for various timepoints post-treatment with control or peptide. Dotted axes indicate skewed scale. (D-E) Boxplots depicting statistically significant percentage change in (D) body column length (n=7-10, p<0.0001) and (E) average tentacle length per polyps (n=min.7, p<0.0006) 10 seconds post-treatment. Significance levels are p<0.0001: ****, p<0.001: ***.

## Discussion

Previous attempts to computationally identify cnidarian neuropeptides relied primarily on screening for known C-terminal cleavage sites, amidation residues [13–16], and N-terminal peptide motifs [17–19]. These were essential in identifying many conserved neuropeptide families with similar patterns but could not cover many highly divergent neuropeptides, which typically undergo rapid evolution through multiple duplications or emerge *de novo*. Through this new pipeline, we identified many new potential neuropeptides by relaxing specific parameters: removing the requirement of amidation and incorporating single-cell data to filter only neuronally expressed candidates. Similarity clustering revealed two clusters of RFamides and Wamides respectively, which are the most conserved metazoan neuropeptide families [10], alongside a cluster of random peptides. The lack of a major cluster of PRXamides could suggest that they are likely highly divergent to the point of overall dissimilarity, or that they are convergently derived from other more distantly related neuropeptides.

We also described the expression of three uncharacterized *Hydra* neuropeptides, with an updated summary of polyp-wide expression in Figure 4A. PRGamide precursor B, while previously reported [12,16], was uncharacterized. Here, we showed that it is indeed neuronally expressed and that this pattern is entirely the opposite of its known Hym-355 counterpart, which is highly expressed in the basal disk, hypostome, and tentacles [28]. This could indicate either complementary or antagonistic functions which are as of yet unknown.

**Figure 4.**
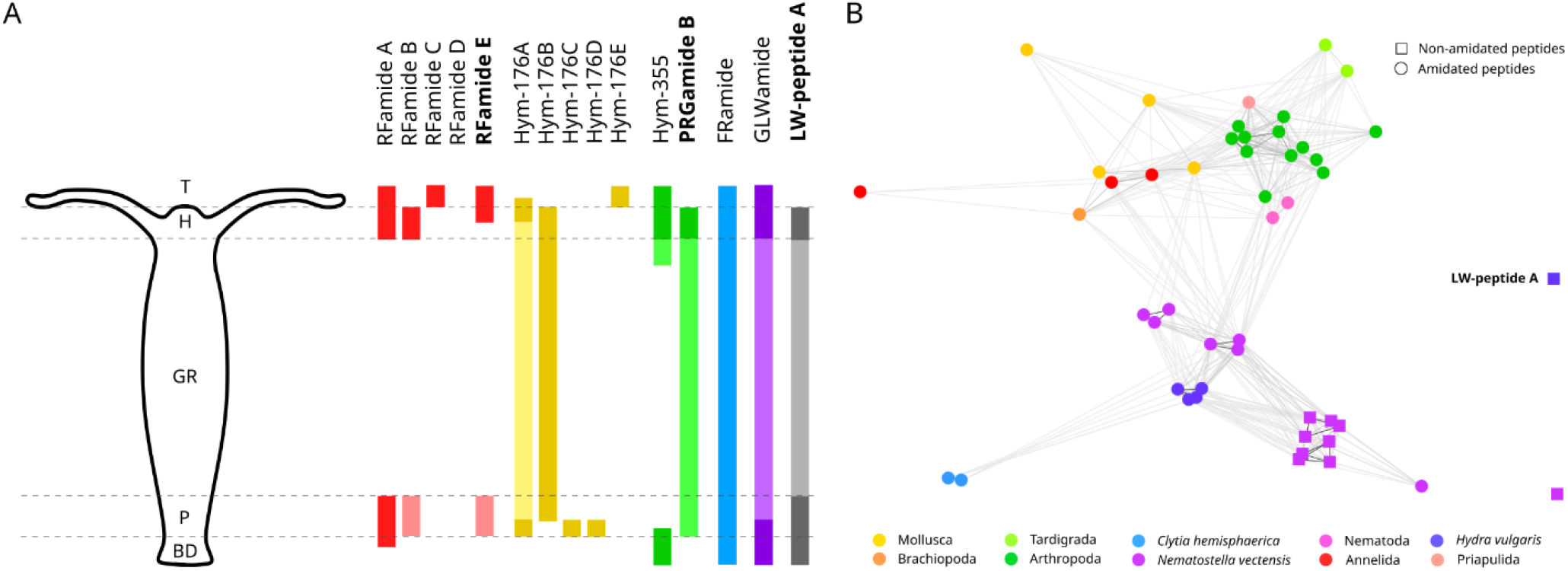
An updated summary of *Hydra* neuropeptides and the relationship between LW-peptide A and other Wamides. (A) A schematic representation of the expression of characterized neuropeptide genes across the polyp. Lighter colors for the same gene indicate relatively weaker expression in that region. T (tentacles), H (hypostome), GR (gastric region), P (peduncle), and BD (basal disk). (B) A network map displaying similarity clustering results of identified cnidarian Wamides and their potential derivatives along with a list of representative metazoan Wamides. G009409 shows no similarity with any GLWamide or related peptide.

*Hydra* contains three well-known RFamide preprohormones A, B, and C [8] and an uncharacterized RFamide paralog [16], which we refer to Preprohormone D for continuity. We identified a new RFamide precursor: Preprohormone E, whose high expression in the tentacular neurons and few peduncular neurons lines up with the patterns of the three known genes [29], suggesting involvement in similar functions such as coordinating feeding behavior [30]. However, unlike the other RFamide genes, Preprohormone D encodes 2 unique peptides, is present at a completely different genomic locus than the known genes (G017220, G017226, G017227, G017228), and does not cluster with the other *Hydra* RFamides (Figure S2), suggesting that it could be the result of an earlier duplication event before the emergence of the four characterized paralogs.

The most striking *Hydra* neuropeptide gene we uncovered was LW-peptide precursor A, which encoded multiple putative peptides with C-terminal LW motifs lacking any amidation residue. Its expression across the polyp is similar to the *Hydra* GLWamide precursor in its ubiquity except for its lack of tentacular expression [31]. *Hydra* GLWamides are multifunctional, with implications in *Hydractinia* metamorphosis, isolated muscular excitability in *A. fuscoviridis*, and the detachment of mature *Hydra* buds [26,32]. Hym-248, a GLWamide, has also been shown to induce body elongation in *Hydra* [26]. It was subsequently concluded that *Hydra* GLWamides are involved in the contraction of circular muscles (elongation) and sphincter muscles (bud detachment), but not longitudinal muscles [26]. These peptides’ amidation and the GLW motif have both been found essential for their specific bioactivity [26,27]. Surprisingly, exogenous addition of LWa1 peptide appeared to induce immediate body and tentacle contraction, suggesting involvement in longitudinal muscle contraction. This also aligns with its expression, being concentrated in the foot and head regions, with a ring of expression around the base of the tentacles. It is highly enriched in the En1N/N4 neuronal cluster in addition to other neuronal subpopulations described in the single-cell atlas [20,33]—this combinatorial expression of neuropeptides in various populations can lead to the generation of unique neural networks (Figure S1). Indeed, LWa1 peptide could be one of the peptides involved in the contractile burst (CB) circuit, similar to Hym-248 in rhythmic potential (RP) circuits, as part of the neuropeptidergic components involved in inducing specific *Hydra* behaviors such as somersaulting [34,35].

To our knowledge, this is the first time a cnidarian non-amidated neuropeptide has been shown to display any significant bioactivity. In general, neuropeptides are amidated for stability [1,36], suggesting that these LW-peptides could be rapidly degraded upon endogenous preparation. Furthermore, since LWa1 peptide appears to affect a separate muscle type, it suggests that GLWamides and LW-peptides have distinct target receptors. Combined with LWa1 peptide’s lack of involvement in bud detachment, a function common to all *Hydra* GLWamides, it suggests complete functional independence between the two.

Initially, we assumed that LW-peptide A belonged to the Wamide superfamily, to which all cnidarian GLWamides likely belong. However, sequence similarity-based clustering of representative metazoan Wamides [37] and relevant cnidarian sequences revealed that LW-peptide A appears completely unrelated to this superfamily (Fig. 4B). This is surprising given that many *Nematostella* precursors encoding similar non-amidated peptides clearly are imbricated with their amidated counterparts. Since this gene also has no homologs in any other organism, its taxon-restricted nature indicates that it either lost all its N-terminal Glycines and diverged significantly to show no conservation following a duplication event or, more likely, that it independently evolved from another unrelated gene, not part of the Wamide superfamily. Its muscular specificity implies different unexchangeable receptors, which would also had to have evolved alongside these neuropeptides or be co-opted from an existing system. Hence, we propose that LW-peptide A is likely a *de novo* innovation in *Hydra*.

In light of such diverse neuropeptides at the base of Metazoa, this lends further evidence to the existence of a sophisticated peptide-driven neuronal communication system. This work adds to the growing understanding of increased complexity in early neuronal systems and peptide-based chemical circuits, with recent ideas such as the chemical brain hypothesis and investigation into peptide-rich ctenophore neural systems providing support to the hypothesis that the earliest nervous systems were peptidergic nerve nets [38, 39]. In particular, there is much still unknown about non-amidated peptides like LW-peptide A and their potential receptors. Indeed, the cumulative lists of peptides and their receptors in Cnidaria suggests a greater neuropeptide diversity in basal metazoans, opening further avenues of study investigating neuropeptide evolution, their physiological roles, and involvement in complex neural circuits.

## Materials and Methods

### Neuropeptide prediction pipeline

Predicted protein sequences were input into SignalP 6.0 [40] under the Euk category and slow-sequential model. Sequences containing signal peptides were input into DeepTMHMM [41] and those with transmembrane regions were removed. Sequences with domains matching the Pfam database were removed using InterProScan [42]. A script was written in Python3 (S1 File) which filtered out sequences with no recognizable cleavage sites within a 20 amino acid range that could yield at least one peptide. BLASTP was performed for output sequences against the UniRef90 database and those with top hits (>30% identity, e-value<0.05) as non-neuropeptide homologs were removed. Hits for neuropeptide precursors or uncharacterized proteins were included. ScRNA-seq data [20,33,44–46] was used to identify only neuronally expressed sequences to include in the final list of candidates. For further classification, CLANS [47] was used to identify BLASTALL clusters within these lists.

### Whole mount *in situ* hybridization (WISH)

Digoxygenin-labelled probes were synthesized by *in vitro* transcription from partial transcripts (S1 Appendix) cloned into the pGEM-T Easy vector system (ProMega) and verified by sequencing. Sequences were deposited in NCBI under accession numbers PQ781191-93. Whole mount *in situ* hybridization of *Hydra* polyps was performed using the protocol described in [48] with hybridization at 55°C. Images were taken after mounting dehydrated polyps in Euparal (Roth) using an inverted microscope (Leica DMi1). Final representative images were stitched and magnified images taken at different foci were stacked with Adobe Photoshop 25.1.

### Neuropeptide behavioral assays

The LWa1 peptide (EPEPPETGLW) was synthesized from Apeptide Systems (>90% purity) and solubilized in H_2_O to make a 10mM stock solution. 24-hour starved polyps were first treated with ddH_2_O for 1 hour to remove the outer glycocalyx and transferred to 792µL of S-medium in 24-well plates (one polyp/well) under microscope lights until they were properly relaxed. 8µL of 10mM stock (test, final 0.1mM) and H_2_O (control) was added near each polyp. Videos were recorded for 2.5-5 minutes post-treatment and snapshots at 15, 30, 45, and 60 minutes were taken in addition to specific timepoints from the video. Body and tentacle lengths were measured using ImageJ software for each polyp at each timepoint. Instances where polyps were oriented such that accurate measurements were not possible were excluded in the final analysis. Statistical significance was calculated using the paired student’s t-test. Plots were made using matplotlib v3.9.2 and seaborn v0.13.2 Python packages. The bud detachment assay was performed as described in [26] and S1 Appendix.

## Acknowledgments

This work was supported by grants from the Science and Engineering Research Board (SERB), Department of Science and Technology (DST), India, Start-up Research Grant (SRG) SRG/20-22/002279 to PCR.

## Contributions

PCR conceived the project. PP performed experiments. PCR and PP performed the data analysis. PCR and PP interpreted the data. PCR and PP wrote the manuscript. PCR supervised the project. All authors read and approved the final manuscript.

