## Supplementary material for "Identifying neuropeptides in *Hydra*: a custom pipeline reveals a non-amidated regulator of muscle contraction and other new members": S1_Appendix

### Supplementary Appendix 1

Sources for genomes and predicted protein sequences

| Organism | Source |
|---|---|
| *Hydra vulgaris* AEP | UCD_HVAEP_1.1, https://research.nhgri.nih.gov/HydraAEP/ |
| *Nematostella vectensis* | jaNemVect1.1 (NCBI RefSeq), https://www.ncbi.nlm.nih.gov/datasets/genome/GCF_932526225.1/ |
| *Clytia hemisphaerica* | clytia_hm2, http://marimba.obs-vlfr.fr/ |

List of Primers Used

| ID | Primer (5' to 3') |
|---|---|
| G0016204_F | CTTGGAAGAACTCGCTTTGGT |
| G0016204_R | AGACAATGGTACGACGTTAGC |
| G004106_F | AGCCGTAGAAAGGAAGCGAT |
| G004106_R | AGCGTGTTTATTTGCCTGGA |
| G009409_F | TGTCACCAGTTGTCCAATCT |
| G009409_R | AGTACTTTTGGTTGCCCCTT |

Bud Detachment Assay Protocol

Polyps with mature buds yet to detach were identified under the microscope and by squirting S-medium using a Pasteur pipette to ensure the buds did not immediately fall off. 3-5 such polyps with multiple buds were transferred to a single well of 990µL of S-medium in a 12-well plate. 10µL of 10mM peptide stock solution and $H_2O$ was added to the test and control wells respectively. After 15, 30, 45, 60, and 120 minutes, the polyps were gently squirted with surrounding S-medium and the number of detached buds were recorded. For each case, three replicates were performed with 3-5 polyps in each well. The proportion of detached buds across various timepoints was then plotted for the control and test conditions.
