## Supplementary material for "Identifying neuropeptides in *Hydra*: a custom pipeline reveals a non-amidated regulator of muscle contraction and other new members": S1_Data

**S1 Data.** Collection of all the final *Hydra vulgaris, Clytia hemisphaerica,* and *Nematostella vectensis* neuropeptide precursor candidates annotated for signal peptides, cleavage sites, and encoded peptides.

**Legend**

Signal Peptide
C-terminal cleavage site
Amidated peptide
Non-amidated peptide

**Neuropeptide Precursor Candidates - *Hydra vulgaris* AEP**

>HVAEP2.T004115.1 - **Hym-355**
MLSLTVATLLLITSIVMAMPNRDATDSNESDILNILDEYIVKVAEMTANEAKILNDVRNYYNDRSSKSLGEFP
QSFLPRGGKRDARPRAGK

>HVAEP2.T004106.1
MMRTAVFGCFILFTIVLALPYRDAFDLFDRFDEYIEKVAKVTADEARLLRDVRNFYKLTKENFVSNADEDD
FQDYAPRGGKRENRPRPGK

>HVAEP5.T009409.1
MTFHLFATFLLIMSPVVQSTRMVNKEMNSASEKELAKFLDEIGLDKEASENRREPEPPETGLWKKENVKR
QHPETGLWRKTSINYVETEPLKRQPEAVLWKKQPEAVLWKKQPEAVLWKKQPEEVLWKKQYKNLKIQN
EDEHHAESKSKIVKQPVLSKRGNQKYLLLENRNFKQTETDKTKEYEEEEASEDENNSILDLVDRKSNEDE
DQEGLWR

>HVAEP9.T016170.1 - **Hym-176B**
MSKVKKLCEFNIILVLYIFLVFSVNALPFKDDEETGIEFDGNISESGNEYQSNQYYDYNKIKNQIYNDYPNIIE
KNFKPLKVMKMGRGANDHFDQIGSRKSNDVNLINGNQQDKPAFLFKGYKPGDQTQKKS

>HVAEP9.T016171.1 - **Hym-176D**
MSKANKLTAFNILLVLNIFVILAVNSLPLRDDEEIDSEIDGDITELENGYQNTQINSYDRHKKQLNPKDKNKK
FMIFQGPKVGRDVDFHSVQSPSNKVGKSTRFYYGNDYR

>HVAEP9.T016173.1 - **Hym-176A**
MSKINKLTMYVFYAFLVLNIYVVLSVNSLPLRDDEDTDEIDGDISELENEYQTNQVYDYNKFKNQADLKVK
SRNHYAPFIFPGPKVGRDVNFHSVLSPSDESRKSFNTYYENGYQHDKPAFLFKGYKPGDQTQKNL

>HVAEP9.T016165.1 - **Hym-176C**
MPKTNKLIRLVFNAFLALNIFVVVSVNAMPFRDNEDTDDKISSDINTLKNESQSSQINDYNKYQKISTIKGRL
QYYPFYNQNPKIGRDASFNSAQDASNNGRMKKLTYIYNKNNYQKDKPLYLFKGYKPGDQTQMHF

>HVAEP9.T017220.1 - **RFamide preprohormone C**
MATNMALLAFVFFATSIFMLTKADQNEDNQKYDGIARSLKVLLQNYNEKQEEKSDIQNIIEKFSEYQNTGK
TIQRKDNVNPMFEKKDAVEQWLGGRFGRVVYDLLLSEVSKDHKRNDETNPMIEKKDADTENRFNREALE
QWFSGRFGLTNHKRNNEANPMIEKKDSDTENRFNRETIEQWLSGRFGLTNHKRNDEVNPMIEKKDSEIE
NRFNREAIEQWLGGRFGRTVYEFLLSETPEKRKK

>HVAEP9.T017220.2 - **RFamide preprohormone C**
MATNMALLAFVFFATSIFMLTKADQNEDNQKYDGIARSLKVLLQNYNEKQEEKSDIQNIIEKFSEYQNTGK
TIQRKDNVNPMFEKKDAVEQWLGGRFGRVVYDLLLSEVSKDHKRNDETNPMIEKKDADTENRFNREALE
QWFSGRFGLTNHKRNNEANPMIEKKDSDTENRFNRETIEQWLSGRFGLTNHKRNDEVNPMIEKKDSEIE
NRFNREAIEQWLGGRFGRTVYEFLLSETPEKRKK

>HVAEP9.T017227.1 - **RFamide preprohormone B**
MCCFHPVEKLKMLSNKKVKLLFALVLIVVEVVKSDDKNFSLEVNKDVKRFIKDILDAKSEEQLMSGRFGKS
LPDEEDIDNEVENEYDNEYDDETESQGIINGRYGRQLLRGRFGRQNDNKAASKESQWLGGRFGKEVAT
QWFNGRFGREIGGRFLPRFGREFNKPHYRGRFGRVAKL

>HVAEP10.T018128.1 - **GLWamide**
MGMFERKKIVLLVSLICVSQQAANVQDANSKSTSTELKVVKPQKRVTPVKDAEKLSILRTQDNSLDLNTNG
EEVWDELTHNIPLEYIEKIYNELNQLAQNENRPKRLWGATAAINTDNLNPEVENELENKKNAPVIEKFERPI
GLWHKDVETKNPENRLPLGLWGKDSEPLPIGLWGKDADVNDDLKKEPLPIGLWGKDIDSTQEDNKPNAY
KGKLPIGLWGKDNALTNDFGKKNNGKDSGPPPGLWGKDSKPIPGLWGKDNGPMTGLWGKKDVGPPPG
LWGKKDQPPIGMWGRAGKKDSNPYPGLWGKKEEEIENVDKEFKEDSLEEYPACLFENPPCEIQEKRYKI
EKSGPPPGLWGKRSEKYSMNKPPWRGGMWGRSEILENSVHDSKQTNTIDMEHAEN

>HVAEP10.T018128.2 - **GLWamide**
MGMFERKKIVLLVSLICVSQQAANVQDANSKSTSTELKVVKPQKRVTPVKDAEKLSILRTQDNSLDLNTNG
EEVWDELTHNIPLEYIEKIYNELNQLAQNENRPKRLWGATAAINTDNLNPEVENELENKKNAPVIEKFERPI
GLWHKDVETKNPENRLPLGLWGKDSEPLPIGLWGKDADVNDDLKKEPLPIGLWGKDIDSTQEDNKPNAY
KGKLPIGLWGKDNALTNDFGKKNNGKDSGPPPGLWGKDSKPIPGLWGKDNGPMTGLWGKKDVGPPPG
LWGKKDQPPIGMWGRAGKKDSNPYPGLWGKKEEEIENVDKEFKEDSLEEYPACLFENPPCEIQEKRYKI
EKSGPPPGLWGKRSEKYSMNKPPWRGGMWGRSEILENSVHDSKQTNTIDMEHAEN

>HVAEP10.T018128.3 - **GLWamide**
MGMFERKKIVLLVSLICVSQQAANVQDANSKSTSTELKVVKPQKRVTPVKDAEKLSILRTQDNSLDLNTNG
EEVWDELTHNIPLEYIEKIYNELNQLAQNENRPKRLWGATAAINTDNLNPEVENELENKKNAPVIEKFERPI
GLWHKDVETKNPENRLPLGLWGKDSEPLPIGLWGKDADVNDDLKKEPLPIGLWGKDIDSTQEDNKPNAY
KGKLPIGLWGKDNALTNDFGKKNNGKDSGPPPGLWGKDSKPIPGLWGKDNGPMTGLWGKKDVGPPPG
LWGKKDQPPIGMWGRAGKKDSNPYPGLWGKKEEEIENVDKEFKEDSLEEYPACLFENPPCEIQEKRYKI
EKSGPPPGLWGKRSEKYSMNKPPWRGGMWGRSEILENSVHDSKQTNTIDMEHAEN

>HVAEP10.T018128.4 - **GLWamide**
MGMFERKKIVLLVSLICVSQQAANVQDANSKSTSTELKVVKPQKRVTPVKDAEKLSILRTQDNSLDLNTNG
EEVWDELTHNIPLEYIEKIYNELNQLAQNENRPKRLWGATAAINTDNLNPEVENELENKKNAPVIEKFERPI
GLWHKDVETKNPENRLPLGLWGKDSEPLPIGLWGKDADVNDDLKKEPLPIGLWGKDIDSTQEDNKPNAY
KGKLPIGLWGKDNALTNDFGKKNNGKDSGPPPGLWGKDSKPIPGLWGKDNGPMTGLWGKKDVGPPPG
LWGKKDQPPIGMWGRAGKKDSNPYPGLWGKKEEEIENVDKEFKEDSLEEYPACLFENPPCEIQEKRYKI
EKSGPPPGLWGKRSEKYSMNKPPWRGGMWGRSEILENSVHDSKQTNTIDMEHAEN

>HVAEP11.T021114.1 - **FRamide**
MYLRLLLVFFVLQISLQESNVRQLDLGQLIEDYLAKENVRREEFLNKINTEILRYIYELENENKGKKRIEASA
DKNVLEKVLTEVPSIRESVTSKESNVNKMHNSLDSKSSIRSIPTGTLIFRGKKESNSNNENASEQGAPGSL
LFRGKKEPNVKENSKNETEASHGERLQQTERNFLVKTKEYIEKLLNSGEEIV

>HVAEP11.T021114.2 - **FRamide**
MYLRLLLVFFVLQISLQESNVRQLDLGQLIEDYLAKENVRREEFLNKINTEILRYIYELENENKGKKRIEASA
DKNVLEKVLTEVPSIRESVTSKESNVNKMHNSLDSKSSIRSIPTGTLIFRGKKESNSNNENASEQGAPGSL
LFRGKKEPNVKENSKNETEASHGERLQQTERNFLVKTKEYIEKLLNSGEEIV

>HVAEP9.T016204.1
MMLLMLAFVIHLINCQNQYTETMSSLGRTRFGKRDYETRESDSPYNTHQKTPFKTFGSQYNPSFRERFN
GDQQDFNQRYLKNSHAHQVKQERSSERNDEVKKSNKHEKLMSRDHAIKNLAQKDNKKVVDGVQNYEK
NGANVVPLSRFGRNIDLISIKKQFF

>HVAEP9.T016204.2
MMLLMLAFVIHLINCQNQYTETMSSLGRTRFGKRDYETRESDSPYNTHQKTPFKTFGSQYNPSFRERFN
GDQQDFNQRYLKNSHAHQVKQERSSERNDEVKKSNKHEKLMSRDHAIKNLAQKDNKKVVDGVQNYEK
NGANVVPLSRFGRNIDLISIKKQFF

>HVAEP13.T025234.1
MQILHWKKMIFIYCLFALTFISCEEDNKVDDYTPPFTCDFGKYMCGYRKCCKSYSDEEKPFKCYDGSYVC
GYKKCCKTVVEKPFKCEYGSYVCGHKKCCTTIVEKPFKCDYGSYVCGHKKCCTTIVKKPFKCDYGSYVC
GHKKCCTTIVEKPFKCEYGNYICGYKKCCPNKYTQ

>HVAEP13.T025234.2
MQILHWKKMIFIYCLFALTFISCEEDNKVDDYTPPFTCDFGKYMCGYRKCCKSYSDEEKPFKCYDGSYVC
GYKKCCKTVVEKPFKCEYGSYVCGHKKCCTTIVEKPFKCDYGSYVCGHKKCCTTIVEKPFKCEYGNYICG
YKKCCPNKYTQ

>HVAEP2.T004963.1
MKFFVQIIVACILASSVQSFGPRINGRDDKLEINRLRDATQLYMESLCFKDEKIPSDMSSFCFCACWFRSD
KASCLSECEKSFGKLSLQ

>HVAEP2.T004995.1
MNIKILVSVIFLVQSVNNFGLGRRFKSNLGKKESDKILSPTMSLVILHKNKILESKTHRAVNDDIDKTYRDDI
NNCFVDCESKNCMRNCLRLAK

>HVAEP4.T008016.1
MPKVVFKQKMLNFLIKQKMVLLSTLFLLWLTNTCSTFSCYSCSAYKKWDCNQVQLLKYCEKGKEQCITIS
YKYEMFININKTTTNTVYKKHCVDKFTSCKSHCRLLEMTGEKNCMASCCNTSGCNKDGYKNSSVSLLSN
VFCTTLSYVTFLVFILILIF

>HVAEP4.T007766.1
MRFMLQILLASILVLIPTIESFAGGRGPGKDEKTETDNLLEVLAHIDSRNYRNRESYYKIENTRELLDQCSVL
CELAKDYEICVKQCKKQFGFYP

>HVAEP7.T012608.1
MNKVHTRLLIIGCLFAHMKCVPIYEVNDEKNDLCKHAYTKKLLYSLGMSADSSLCILQDKRDVESLFTSVLE
TIFLRQVRKEPETDSTIIESREYCCSPKCDSFMEKLGKPHVKLVKCT

>HVAEP6.T011508.1
MRKIILLWFLVSILSFNNICCLTLQGQVHRVHTGKREGDYKSLYKLVNAGRNNYWKNKRSMLRLQTLTND

>HVAEP6.T011508.2
MRKIILLWFLVSILSFNNICCLTLQGQVHRVHTGKREGDYKSLYKLVNAGRNNYWKNKRSMLRLQTLTND

>HVAEP6.T011172.1
MKMTMVQFYAYLCAAFTIKVICTKSQRKPLEQNAKDVNYNKLITLVTEVEKAQKLAFSVLKSLNNNILSKNN
SLKSNIVSQQKNSNDVILSKVISPVNANGSKTETLYTVLRNKMVYKIKKNDMKSKKKKIYGFWLG

>HVAEP2.T003594.1
MYRWIIILVLLDFVMSKKECKPEIPPGLHDQLEIKNFILGDSKNMKQNPYFTKRLVSKKKESLPCKLWGVEC
I

>HVAEP2.T004141.1
MITLVLFCLFIFVSKHSLSNVVFIIREESINNSKEKKAQEKEYKERKKREETNLFYRCKEECVSDRSIFKAEK
PKQCPLCQIIMKLVCDKQKCIVDGNRPIMILPKFTTEKITTVKKLFRKSYDEKESIKDSQ

>HVAEP1.T000427.2
MGYVRCSTSNSFIVCLSLCLILSIVSCASIENVNPADYIYILKGTGIKLEWRTTYNLFETLMIFYCGYQENHVL
VPLAFDVFGDQFDTVFTRTPPKRYNLPDYNNTWVKKIDGYYDNTLKTYNLELLSIDRNITCGCKLRYMNN
NEEFLITKQNSALVLKRHQLYAQKTSFLDEIRVG

>HVAEP2.T004997.1
MIQIVNLVMAGLLLAHSLDGFAGGMVDNFRKKEDIYQNNKLIEWKRYANLQSNDNKPNVLNNIRFPREHF
KDCAKYCDNNDNNCKNDCKRIAWLVI

>HVAEP3.T005478.1
MQTKVTFVLLLVCIAVSYINCIVAGGNNMHGDYVKSLKNKLSLDGKVACECCKICQNERSYARKVEEFEN
EENL

>HVAEP3.T005951.1
MSSYICLVFLVLSTVTVFYSSYASSQETCGLRWRGCKKSIQNNKAQNTAWLQKNKNPKFALVEKELSNLI
DMSEDEEMNENAKILKRKNKT

>HVAEP3.T005327.1
MLRMKFVLLIFTLAYVSSKKMENQFLCHSCTSYYNLCQPTVSEPITWHLFKKLNRCMQEYVDCVSKFCEE
NLTNTKKMLKAKQLFHLKIKSLQLVI

>HVAEP3.T005327.2
MLRMKFVLLIFTLAYVSSKKMENQFLCHSCTSYYNLCQPTVSEPITWHLFKKLNRCMQEYVDCVSKFCEE
NLTNTKKMLKAKQLFHLKIKSLQLVI

>HVAEP7.T012652.2
MINIKNISLLMAIRCWPVLCFTLFFSNIQSESFSITRKRTSNSDNEDQFSNELKYRDLKEEYEDSVLKRNIRN
ESKSLDVKTVKSYLELIIKDLRRHINHLKSLTKLKDVEVKMQDNIQEGNKGMNCNEPNMEDDNVNKIKDKN
KGCKFCKVISASKYTSSNDSTNYKNVTINENSTIHVITNNNIQNKIITVNKPTAENASNSIKNDRTSFVKNLTL
VEIKTQLIYFNKSKESNVRVEKMANNQTKSFSIDSAEVDKLIQLLTESMILDITESVNKRAWDAVDDEEEEE
NDGSKTVQESLNKLTEKYLMKKRSDWIALLKKKWLKKPRVKSRIKNKKNTKEHTQDK

>HVAEP5.T010060.1

MNTLFFLFALWFFIVCECFQLHDIETDKTKNSYEVEQDWTNIDYRPWREGDVRSNTNRAQVIKKRGRPW
REFHSENEGFQNDQYQKDFELLRERRLASNPLKDVRKAKVHRRPLTKRREMNESLEIGEHRDHKKHLGS
INVN

>HVAEP8.T014819.1
MQMFSFCIDKCIELIIFMKTMSFFVLGFLYFSVYSCFDCQNGPPGKYCSDNLSGYYNCLNNGTSTYIDCQP
GTRCSCYINNPCQTKNICSNYSIPSSMISSFVLHYKGFQETQHPQGTYIEDLHGTIRQETAKKQYFQENIV
GSTHTFVLIIPSKKEFKMYKGTVNRSCSGFELRKFDVFLNELTYFTKISKKVVDNYTNEETFFFRNGRRHL
GQSLTTWEWVVKNNFIKKTLEPIYFKSQFYGGELARQIKITSWTSISIVPSYSNDTFFQIPKICKK

>HVAEP8.T015083.1
MIKALIVIFVLINSVFTRCNYMRWSGCEDSEEAGKSNQEAMSKLGLLFQSKFSGKKDPEVKFADQREDLD
SEKFNDELNDVLRSNLVNFDEKKPDEKNEGDVNHITEETTLSEKNYKIFKKMDNIIHSLMGIPLRIDQFVAK
NRRK

>HVAEP8.T014897.1
MTITILQAVVFLNLIIFSLTATNCEYFRWSGCFEPEVSKDKEINEFVAKIGEFFQHKFQGKTLVDKKKSPINKI
IKADNNESKS

>HVAEP9.T016811.1
MRLGFLVLMVSNVTSFQQNPINKNTELIDEIKEYPNVDNNLRPPKLGRDLYNRGSIDKQNYEINQFKEAIRI
KLRSLVKNSNNKKRSIESRSLVKKCIHA

>HVAEP9.T016549.1
MMRLKFMIAFAISSNVICIASRYQNQKNPVDVSRESFTAARKTILQLCSKDPLCKSQYFDSPAVPNAFHST
KRRLKSKYQNILTYRLLMDLFPE

>HVAEP9.T017554.1
MAVNGIIILCILAFCACTSAMVHGDYTCLALCDPPYEECFKQINKVEDTYFVCMVERSKCVKRCEKFKLISK
ESDSLDQ

>HVAEP9.T016345.1
MEFKKSFFVAIIASYFIVLSFAGRITYDKCLSHNDCPKGFCCAYAHRISQTNGFCLKNKEIGETCSISFLEDD
VFTCGCDYGLTCTIPSPDFKTKRNKSSLAEPKKYRCMRIPEETKEEVELNTNV

>HVAEP9.T016345.2
MEFKKSFFVAIIASYFIVLSFAGRITYDKCLSHNDCPKGFCCAYAHRISQTNGFCLKNKEIGETCSISFLEDD
VFTCGCDYGLTCTIPSPDFKTKRNKSSLAEPKKYRCMRIPEETKEEVELNTNV

>HVAEP12.T021930.1
MKRIITVLLLIQLFSFIFATFQAFNCEKDCPINQCTGAVGECFKRRFACKYICSEENEIENKEIDTKDEYLPLK
EDECYDK

>HVAEP11.T019444.1
MGKIVFSVTLFCILSLYNVFGYREMTKRFFCKPGECGPGQCCKKYFLPPASVCMSYIPEGERCMFGDAV
NLGCGCAPGLECMFKKFFWYCVKPNNEPEPDEEEKEINQLGNEQTNEDDLNDERKYFKQDK

>HVAEP11.T019444.2

MFQGSSYPLVIFEASQTNSSTKSISLKKNMGKIVFSVTLFCILSLYNVFGYREMTKRFFCKPGECGPGQCC
KKYFLPPASVCMSYIPEGERCMFGDAVNLGCGCAPGLECMFKKFFWYCVKPNNEPEPDEEEKEINQLG
NEQTNEDDLNDERKYFKQDK

>HVAEP11.T020256.1
MMLVCFIFIKLMFLFKLEKALGLHCYHCSATNYKTCNESSIIKECPYSSQCVTLSVEMYRKSWNVSTLSQIY
KKSCVVSKYDSCEKNCRFYKNIGNFCVANCCPTNLCNSKEIVIPSFIHRRSRSSAICNSSVCLIDLLIVVFLF
FVKEKV

>HVAEP12.T021928.1
MKIDVILLIAIIAFSPIYSTEDGSKNEFSILEKDRCYFQCKVKKIHCQHVLKLFRPCTKDYRSCAKNCR

>HVAEP11.T020678.1
MSGFIFLCLLLISLSPAMNSSTDDSDINEESSDKNDVTEKELLSNIQSFLAKDFNLRSNSMYSYELLEMKQT
NKKIINDGMSIHFMLTFAQTGCRKNGALKEQLKCPTGVLMDKNKPTKTEIVKVIVNIPEKVLHNFEERHLTVI
YS

>HVAEP12.T023803.1
MKIQISFFVAFILLSSLMAVPLVNKKNSPLSGESKILSGILLSENYVSKRKRKSLNIKKEPIFGFGCYTIECKM
KEELRIILIDKENKDTSNEIDIQFDKQSKFFKTRVV

>HVAEP12.T022979.1
MKRAIGLTLIFIFTTIVYENEAFITSERWRKKPMVDKTEKNPINIRTELCQTNYQWCREYCRNQLNDLSIETC
IARMKSIGFTKKYLYVYT

>HVAEP12.T022883.1
MIHIFKICSVLCLITLVVAHFKIGKRTSYDTSLYKKKISSQLLAFPKTREEKLRQCKKMLTSETNLMNEEDGIS
KLNERITNRRNLILDDNNEKIKESSWKKTIEVICFHFLFSKDNSTKK

>HVAEP12.T023387.1
MKAIILLLILAVFVNDVCNLASGNWVKKEKSQESTLYHIANEVKKVLTLIKGMLYENRQRDSSLSENEEAVY

>HVAEP12.T022859.1
LLLTFYSLTFAKGHYKTNDKSIDCVAQQVESVFDKASIPTKTMYRVVQLINGYHDSYRNLMKSFQEKYNSL
QGLYFDGRKDKTLVVEKIKLKRFR

>HVAEP14.T026486.1
MMISGILLVVFLNTIELNSQRKAIKSTCGYFNFKLCIEEKLMIVKNSDPEKTGFAKNPYSMRLDTLRLDTMRL
DSEISEQNKNLKISSHKKKKNSVKTKNGTYTLKV

>HVAEP14.T026484.1
MIAIILVAIILNYSNVLPQRNSLKSTCAYFDFKMCMKTKIMVEKNVVYDTLAKLDSEVKRKKFKISVPKCAKE
KFGKWRCFVEEKLV

>HVAEP13.T025675.1
MQRNSYLKAILILISISLCSALKCYTCTSRSEASCKVTQKFEECGTNENTCITVHHTFLIYSGNKTMVKSRFL
KSCAETSRNCKAKYCKRFQECKAYCCHDEGCNEFMNPVVTNAKHKSFENSVIISAKRTSLGKKILPFFSLI
SFLVVLQYL

>HVAEP13.T025622.1
MLFFLLSIYVVRCGAEYKGSGAIYMKRNHQDDQISLGTLNKLTKLQCVHACIKNKRCSSFIFDKKDIYLGDC
VLQEKNKNENIQTPIFQIEHSLQNYEIFESFSLIYSSCQEGYENGNRLSGNYFLSNELNWHFCNMGELKNC
GVGGWTLAMTIDGKKPTFAWDSPYWLNDAIYNVEKGFLSPFLEEAKYPAFSYLEFSEICLGMKFDTSEISY
MKIARSGKDTLKRIFNAKFTNIENGRESWLKLLNGSKLQPDCSGTNNEGFNIQGKAANLRLGITCYDDNPE
WSDSWVGLGGQDLFCLETESMQAQSAAGNQCCSNECNEFSLKAMAYLFIK

>HVAEP14.T025882.1
MMIGKFSLVILLLFLLDISDSKPQGCISIRYRRACTNNIGQRSDAMKRQRDEYDGSTKKVKDDILLDLYNVF
NEPS

>HVAEP14.T026552.1
MTWIFLLYIIIAGQQGGQDVEAVGDELESDWSVDLFPMGADGPSGRVQPSILSFFVPILHNLQRQLNAGQ
GLAEQELSVTLTFPMDTDRDEMTVGTKLPVDVPNSKEQLLKDVSLQKLLVNLLQLFNYRSRATSEPSLGL
VQIKPHNHLPDFEKKDVLKLKQSIKRAAETTSGSLRDIFDQETASNQSPHHVSFGMMESTMYCRRRSVQ
SQLPANWGELDNSLRVNNISKFLDGTDFYKGYVRINNDYAHIFIGKTVVASPTDLHVDGTLKHASSIQICYR
VKFLSDLYSIERSSSFSLLSVVKKSGKKVISSKKTKNKDDDGLCFYITEFPITTNKRLPSLKHPNRRLRCKE
KKCKNGNTKRKLTAKSKAESEKLEDALNKYLKIGNANKESDEKNEKESEENTGEGTERKKKIN

>HVAEP15.T028625.1
MQIIIHSMAIVFTTFLTFMSSTSQATGFQSSCRIPCKNIFVNCIHSNTAETLEKMFSHCQNATKSCFLLCYYI
SKQKRDEMYNKVKNLRTIMAKNHYLEM

**Neuropeptide Precursor Candidates - *Clytia hemisphaerica***

>TCONS_00006521-protein
LVVSISEKEKNQFSNIVSKGNMNCVLVFLVLFLANNVYSASLTREEDALVTKLLDTIEKRDAVPRLGKREVP
RLGREIEEVPRLGREIELVPRLGREAEEVPRLGREVELVPRLGREVEVPRLGREVPRLGREMEEVPRLGR
EIDEVPRLGREVPRLGRELEVPRLGREVPRLGKREIPRLGKREVPRLGREASTYDLKQLYNQLKREVSND
MIEAEIKEEKRVLKAFNLGTRGLILRRIGDKLNKDGFSDQKRGMNEKESSRPFLHVKRTNLLSLIEKLTSEE

>TCONS_00068562-protein
MERKFLACLFLLIVLLNLNDGKNIAILIEPDDNLASELEWLGSDMTDSHSLNAGAWPRPGDARSSHDAWP
RPGKREFYGNEMFEKRPFPGQQMQFSWPRPGKKETKEDTWPRPGKRESYSEGDMDSRSGALRRSEE
KETNEDEKLENAWPRPGKREFYASRKMDVRPRGGRDSKSHKISKRNSEAISYDEIDMMLREEAWPRPG
KRDYHMLSVTRPRGGKDARPRGGKDSLRPRGGKDAKPRGGKDSVRPRGGKKDSWPRPGKDAFVNEIN
GSRPRGGKDASKWPRPGKKDIK

>TCONS_00015710-protein
MLSSETTIRILCFFIAVGFAVGSSSPEEEGQLLHVKRETWLNPGFDSMLHRRESQELLNRPRPGRRELFD
LMNQDSILKKRALLHRPRPGRRELFRNPGLDSMLHKRGQEFLSGPRPGREIRPRPGRRERHPDSMLHR
RSSDTDLDYQHLLRNPRPGRRELFRPRPGRRELQHPDSLLHRRSEEMWSRPRPGKREVYYENDGRSE
DEKLLRVLDELKRDIIDELWDRFQN

>TCONS_00015647-protein
MRLVNVVLTILTSKIALGEGRPPSRESDIDQQLIRLAHEIRREIQPGPFGNYRGYAPPRLGKRENIDGKKGV
TSQKRRTVEALDKIGQILLNHRKSLSSDSEVFKNGKKMMTDHLDTTSRKRTHHFIKAKNVPFWYFNKRGL
SNQVDGKKTLGDGNKPAFTHHSQPTFDKFADDVIEKLTTLNKQIKHDAINERDEKMKKRERVGMKSKWN
FWPPRLGRKRETTLNRNMEKAKHPYDRILDDKKGL

>TCONS_00007179-protein
KYSKEVMRRSIILFLSILCTLQFCHCFSTNRFTEIKAWESLREIEPPPLHESDEDFPPALHAKRLASAFRPRP
GKRSNLEELRDLVSRELETLSNLEKENDNFARQETLPPQFQYYWRQGKRAEKPRYIGIRMGK

>TCONS_00025986-protein
MQWFDTVFFSSIILLLLLNFVIAEQHNHEMTPPPGLWGREMPMSMMRVQRQKEKKLPMKFGRETSAINH
ALPMKFGREATTRDRKLPMKFGREASTRDRKLPMKFGREATTRDRKLRMKFGREATTKDRKLQMKFGR
ELPMKFGRSSKSEHQEVTTFDRQLPRIILGRELTTTTDNQLSRRKSTKNLLKRRESFVEQILSNEIRREFFN
KDIVKRFLTLYWNKARQDRYKNNKFQ

>TCONS_00029927-protein
LQEQNMKIYFGCLFVILSVNQIGCYPSSNQNSERELVRRIYKTVHPNPHYQVNEIQRVKEALKRRVLENVH
RVDSLKASLKRVLGQDAGNGFHMSSSKIFQKKRSKARLPHSYMFRKRQNSPGALGLWGREVEAPGDIG
PPGIWGDVVPDETRKDKPGAVQGLWGKDERVIRALLKTLKR

>TCONS_00069868-protein
MDSTHLCTLLTLGVVFFSTTSAQPYLRPSYPLKYQRSIPYEPGYEANPIRWGKRNSLYQDTEDAGLWGK
RQEELGEGLWGKRDIDDVRLYSKRVQLKPRQDPEIMTTNDISDLLERDNKAMVESLVKLRLDVDEALRHF
NIVEIQEDKRQDLESAGLWG

>TCONS_00033583-protein
MELKYFLASSIFVIIAIQLASCSSKAEEYKQMKKEVDGLLKEIVSQENAKQHTSEKKSSQWLNGRFGKRQL
VSGRFGRELKQWLNGRFGREATEQWLNGRFGKREADQWLNGRFGREVEQWLNGRFGRDAAEQWLN
GRFGKRSANQWLNGRFGKRSADQWLNGRFGKRSADQWLNGRFGREASADQWLNGRFGREAADQWL
NGRFGREAKEQWLNGRFGREVEQWLNGRFGREAGQWLNGRFGREMGQWLNGRFGREADQWLNGR
FGREADQWLNGRFGRDAAPLAARYGDEPAHVESQTIASPEEAKPKVVAAVKVVKKPVAVSE

>TCONS_00055959-protein
MLLILAFWMVLLQQVSAACYNGQVEGSIISLRNGETSECKQGIATKLSAWRNTKREDVTASDTEKNQWHF
ELFGDSRPLSFANHRGPWMRAGRKDEPRPWLQFGRRDINFEVFHSDKYNEPPFAMINGKRTANSKKIYS
EKINELPFAIMKGKRKVNSKRLHSVEINDLPFVMMNGKRHARDNIVGPKGMSLQWASRFGKKDNSKKEL
KIKNSNSKRNTKPWFDIGSEGSSLQWQVNPFGKRGKRDTLPWYELEGGGLHWNRRIASKRSQRLQLPP
RVPHKRGERY

>TCONS_00019380-protein
MKNLLLFCVIANLVGFSSSSSFLYDELEQYLENEMSKRQNIFRWGEGKFMNGKRSTPLDPFTISKRQNILR
YGKGQGAFAFGKRKVPLDSFDISKRQNILRYGKGQGAFAFGKRRVPLDSFDTSKRQNMPRFGKGYGVF
TFGKRSVPLDPFTISKRQNILRYGKGQGAFAFGKKETNNFKRYLRRFWNTDEFIKALLKSQH

>TCONS_00007166-protein
SQCRHNKNNKTMRSLLAVLCLSFIVCEAASIVKKDASVKKEVQPEEDIEPAVIYTEAVVDENGNVLEKDTII
KGSKGRMFINKDTYDKETRKRSTDTLVQGSDGKASIYSTDINGKKELVLVEEPDGSIDAVDEKELEEALEE
EEEEEEAGMKKKREKSGPGILLRGPGSRFMFGRKREGPGILLRGPGSRFLFGRMVEGEEKTKKEAGPGI
LLRGPGSRFLFGRDQKEKKEKSNEESSKEKKEAGPGILLRGPGSRFLFGRDAKESQKKEAGPGILLRGP
GSRFLFGRDASKDTTNAELKTLIKKALSEAKEKQSKSTTKKSE

>TCONS_00019378-protein

MAKLTGLTCLVVIAAFVLKCSTSPVPKDLYKGEAISKRQNIMRWGNGNFAVGKKEVPANGKEISKRQNIM
RWGNGNFAVGKKEVPRNGKEISKRQDIIRWGTGNFAVGKKEVPANGKEISKRQDIIRWGTGNFAVGKKE
VPKKQ

>TCONS_00047380-protein
MNSSWNTQFLMAAVLIGLVMAEVRTDETEALPMYLLSQQLYLKKSNDEMSKRSELPVGLINVKHFGKEN
GYAKKDIPMRFLSKKDIPMRFLSKKDIPMRFLSKRENEEDIPMRFLSKRGELPMRFLAKKDDEKNSEEELP
MRFLQKRSQEKLCKRVVDICSNVLDQKR

>TCONS_00049488-protein
MELHLLVSFVLVLIYPVESRRSFRYGKLLKKQYDGILEDMKQHTDLVRLQSITKRGALPQGSLRSICQSFCL
SNDCYGPPCDAFVYGREERRSIKRGALPQGRLRSVCRQACRSENCLGPPCDMFGYGRETEREIKRTFV
GMSQGISIRKLCRQKCANNNCFGPPCNVFGYGQEVKRLVNSGKESGIGKRRMETIWNRSTFAQGIVDLR
SICREACRNRGCYGPPCNRFVGKRSAGVYGDNNQVEKRVSFPGSSPLKVHCRLCNLKNVECLPSGPCR
KFRNRKSAE

>TCONS_00058018-protein
MKIISLFFASFCINAVYSKSLNDKDEDQQSHGNMVGKGPFRIPHKRESYGMIVKKENEAEDHTWKESTKQ
NGLQKRKVETSGWLDMKKKKRSGDEEDFQKPGIDEVKDKAFIPMKLYALSRKRSFAAENNPISKWYKR
DSENGNPIHKWYKSEVRNADDDVNGEIFEDKLEQDRLPTRWMDELSRSGTNQDLEDEYEIKERNFKEDD
ERKMRQNGLQKKDDYGQYDVEYEKIMRRQADEQQKKNDDYRRYNEENEKIMRRQADEQQKKNDDYR
RYNEENEKIMRRQADEQQKKNDDYRSYNEENEKIMRRQADEQRKKNDYYRRYNEENEKIMRRQADEQ
RKKNNDYRRYNEEYENLMRRQADEQQNENLPTRWMKDEYDRIMPATFTLRQHPDTFIPKREHVTGYFD
DDDAITANGNRRDSRRFRK

>TCONS_00041248-protein
MXLTAVPLACLLVATLSNGLSIDGKELANQKRGRYSYSPYPPVNAYIRRLSMDYLRRIKFLLASGRKASK
RSASDDALTNAEDYLLKRQDDMKTIMWGREASPMLSREADTPMLTRSEDSPMLSRDVPDFERDLNEMF
RREFGSPMLSKKSSPMLSKRSVENIVQENKAPMLSKKSSPMLSKKTTKN

>TCONS_00001513-protein
MTGKFVAAIFIFGVLKKADAVKCYTCSTLGDPQVCKLWQEMEDNQRECVSGSRCLTISYKESHNGGQRP
RFVKKCSNPTLDVCDSHCKNVKNVIPNTCRTSCCSTNLCNEEVNEKYIWYAFNTGKSRSNSANTKHMKN
DLVTFCLTSV

>TCONS_00001406-protein
MDKIQEYFKILFWLTFITDSLSIECYECSGTHTYCNNSRKTTTCDGPQYSCLLSSSTTRVQVIKNGALGNEI
LTSVTKSCGVETQSSYCKDKQDMDPNMLLCQTNWCYKDFCNFDLKDIKKGAKIQTNQVDPYGWHKTSG
AAPIKKYTVILVWSLLVVLLLIYF

>TCONS_00001514-protein
MNRILFFLFFLATNLCYKEVSSLKCYTCSTLDAEVCRLEQALDESQLECGARTTCVTMTYEQFSSGQKIKR
FIKKCGMADSRRCKDNCENVEHLVKDSCKTSCCSSNLCNEEVKEDIIWYALIGTPNSSHRIFQGNFFTIFA
VIVSGILYXFQGNFFTIFAVIVLGILYLV

>TCONS_00002273-protein
MSTILEGFCISIFLVFTCVKGTLSIECFDCSGKIEYCNKTETTKTCHGSQFSCLLRFSSTKVWVIENGRLVSE
IQERVSKTCAVESRLYCEDQKKIDPNMDYCKAHWCYKDRCNFELKQAGSDTLTNNKIDPYGWYRTTSSA
RKNHSETKALTVWLVCICNAIFIRNIGDTVR

>TCONS_00002660-protein
MNRQILIFLTLVTCVVLVKSRSSRFGAKKREWPIKLCTADEDCEGGCCKGLFCFRYLTEDERCLFGHHLK
VGCGCEAGLSCERKKFSYSCQKPEDEDESEDNFNRELEEILMRAETRRAVADEGENADNASIDNETKRN
VSQF

>TCONS_00005292-protein
MKNLIFFVTLSSLMAGCFSVKTKNSDSDSVKNNLKRIINTALDALVNIEDNANGVTARSDELSAQAAATLSD
EETARKEIEEDDSLSNEIKDLEKILDAAEKRDNPKRESRDDKGEK

>TCONS_00008023-protein
YLSTMEWYQVLILLVCVQCAILFPRQQQQFRNGDERELYIKVKRSSFCNRHFTICGWQRGKRVSETIDDQ
NEIRSIESRNNQLEKRSRYFLSYCRQKCFHLTTWNDFRDCVNQC

>TCONS_00012798-protein
MCSNNFMKLFLLLIFQQICALESINFESFQVDEGECSMGEPGIEVKSTTECLLHCGMEKCMNSLIKNQTCY
CTDTECVPQRKQQSLHEANFFTSPLQKVTIEPICYGTKGSSFASFNVPQSGRIKYFKLIHVSGYVSCSNVN
GNSTWACKVMEPSTVLTGITNTNNQIIAPPGIHDNGKFQIQGVNHMNDRELILPAKSSIEVKKKEELRIWYT
EDLVDYKEQDNIGTHCIHVLLKYC

>TCONS_00014019-protein
PFSVCRFQWVRQSWRHILKILNLNFYYRKSIQNTEQIMKVAVVLTIFAVMALHQWGAESASCQNDCTDVC
LHDTNVNNPFAYINCNKRCKAGCGIVNKRSDEYASGLLLRRALTQRRMAEREW

>TCONS_00018205-protein
MSWICVLACLTMVPGWSSTMSVSNKELTELETLIKAYKDISGNIQHRAVSNPEPVIPKSTNGNSQLHEQQ
VEASKVGLKIMAASHAKACGTASRIAFNECNEKANTKNEEQHCAETYVDKYSHCYFGETLTTGSSDLMH
CSGSCMWNFDNCLVNSDKVEMFICMNGRDICSNNCPWSNAISSNSKRSGTNCNSVCEGKFDMCFNSA
QKGSHIFLCNVSRALCRKQTTCVIDK

>TCONS_00025987-protein
MKSVACLLLIALIVVEGRSLNPLRSIKSWGDSGTQKGLWGDQMQDQERRQAPAGLWVGDETSIPHFPED
EDTRIPLFLNDARPHLSGISEWRKRQVPLFYNDHIPSFFHDEMSDEDEESTEKRQAVPVYYNKDMGHHY
GIWKKEILNAIAKELKAKDSSPVKDQMX

>TCONS_00026344-protein
ELETSKPQDLFKQHILTMAKMASKIVLSVVMVCLTMEFVVARNENLCDYNLFQDNQVVTAEINGEYDGET
TMLLNYCSGSFRSCGEISHFGQRNRCCTFAKNFCNQQPACGDYCACWREKCRGK

>TCONS_00027160-protein
SISFLCLLVDTVLGLICQKCTSTSIRNCDRHIQNMTCIYPMDICITMTTSYTIRKQNGIRETISEVTKRCALSK
YGCENMCNVQPGHNCQIGCCTGNFCNRTPDPRILRSINNSINLREHKLFRILKILFTCSFVIMYLQ

>TCONS_00030208-protein
MWYARGLVYLAVISVSLSCVSGIASLANAALKKNKGSTKTEFKNFCINFKEFCKFMYDAPASRSDEKTDIK
STDEVGEETQGEEGWKKYRDNSVEEILYNAHSTLHSGSPSRWRLTGQQKDNQKWKRESAFFRKYNNL
YRILLQEQYKRILSSGDAEKLYRRQTVRSEDKNNNMNQVMAFLNRNRNYAIRKDKLVETTNTKRLNRARF
SPEDYFISPKTNNYFVDQISNHLSPTLNKKRHYFQSNVNSRIKRILDNPDFYPNGNVNDVLGKKRLLSAIQ
QVIETSQSEKDSIKDDKMNDILRSTLP

>TCONS_00030260-protein
MKKFSIGFLLLVASLSIAQSAKEEEKGQPSNIESSNEVNGNVPNRPSEEIYNNLPKQKGEHKKCRDLPKDQ
RKKCFQTMVAERKAHQIQQRLQRNGRRYDRIAKNLEAANNEEKAAKIKARMEALGEKNFNLSGKLQEITE
KMKARMKNKKMKKAKAKNVTEKVDKSQGDKNPDKKGKKNRPASKRPAHKKPKEENKEEKEGKNSKAK
ERKKNAAKKAKKNQKRKRNKNRKKGKDGKKEKEQ

>TCONS_00030302-protein
MTQILMTLALACLIHQTTGYLNLLGKLVESTGFPITLHCPNDPGFVLVGGITVNNEPLEEKLQQKVKDQCD
RQMKKSKGSNGGQGATCSLNLPRVNVSDSDTVRVVYNCHDAEPTGNEKWGEEEKKPKIAEEKEEEAGI
DMDLLLACQRRNGVDDVITIERIYLNGNEPPEGEHPDLVPQKMTDKCYGEWTNELENGQSKKYYPVCNI
VNTYGTTLKVVYICLGPGSKRKKSGNGEGNENGGGNGKGNNGNGNGNGGGNGNGNNGNGNGNGGG
NGKENNGNGGGNGNGHGQYGQ

>TCONS_00032825-protein
MLSIVCSLCLFSLIFQVEADAVIRKGGQWEEWRCGKCGPKVDPSKYVACKRRCFVNGDVVPDYHCDGW
GEKFVKCAPNNRDTTKACMECKWPTPCSDDQAFGDLEGICRNPIVQKTRCGNIPSNNGFYIACASPAWC
MPCATPQCLHGTF

>TCONS_00046937-protein
MKFNISIIWFWYSITLMSKYGHSSTYEGYHCYQCKAEDEQQCEKSFKTKSCKLGCAITSYRFRSVFSENR
VIEETRFEKKCWKANDKDFCDHLKIVNKGVTSCFARGCKLNQCNWNIKSLTDIADYGRYRMNSSSRINNH
HHLICLIIMTFLFVSLYILVFSIK

>TCONS_00050932-protein
MPPFAIYLLISHALLHLHYAQPTKADIDGKDNEKHVTANPKETLENSVIPLLEEEEKEEIGAQNPTNAQQNA
QRKHRQVIIKRVKLAALAKDIPTAKEGELDCEMILSEECFGELCFRGMCTERCIPVERKVCKD

>TCONS_00055425-protein
MNTKVIVSCILVVILANCQAHFRIGKRSKFVSRQRQAAPPHNEILNFLQKAGGSRKICQHLLRSSNSGLEK
NEAGSYVIDKLQSDVQALKSSHTSAITKYKALLELLEEYCLQNALNAALTSNEDASQSEIEDEVYRK

>TCONS_00060728-protein
MMISPRLFICLVFLATAYSACLNDREADTTIGAGNVKKAFNEDQEQIARVVSQTNVENSLINNSGKLVNKD
VKSASKANIQSPGKKESTNIDEEDSSVLTKDELSKKGDCMCIYNNLICGNHRVGFCQDRRHGRKRDSELT
KKGDCMCMYNNLICGNRRVGFCQDRRHGRKRDSELTKKGDCMCMYNNLICGNRRVGFCQDRRHGRK
RDSELTKKGDCMCMYNNLICGNRRVGFCQDRRHGRKRDSELTKKGDCMCMYNNLICGNRRVGFCQHR
RHGRKRDSELTKKSGGCVCIYNNLICGGQMLGFCQDRRNGRRTADSAGNDIGEFKRSDCMCFHNELLC
GGHRIGFCQEH

>TCONS_00062707-protein
MLDFMRILFCLGFVVVAVIAEAETQDDNLAKRVSHQTHIPGDKSPGRKFILHLRPHGDEQTKQRKMLPLK
RENTAVPAGDDLFASRFGKEDEDAAESTWSAWRMGREADTVDTTKRVKDAESAWSVWRMGRETDGT
TTVKDATTRERDASIPSHPRYQVFRSGREAEKSLPMSRAFGGKRDSPGDAPIPYYYSFNNEGKDFSWNI
FGRKRAAESDLPMSRFVDGRKRDLPMGRTFSKRESNRMFSRADVPFFVKRGSNLPMNRMFSRSAVPY
FGEGKRESNHLPMGIFSREKTNADVPFYVKRASSLPLGHMFSKRANFLNNRLPMQLGFKRASNLPMTRM
FSKRASLPMLLGVVKKEHGGFARAPEYNWGPASLKYHRRSLLPMIMHERKRRFNGFRPVVSGIINNDRN
RKLKNNVNDFHVKKL

>TCONS_00066213-protein
MNLFLVISLLFGYCFISVNCLSLASIAECQDIAPSELCHDCIRYSHVCRRSCHVCSAHDEETNGAEENRLDI
FRREILKRLRNDRISRILRRDEDELENAPSKRRDLPARYACSITGDCHKT

>TCONS_00066253-protein
PSFITMRLSSILLIVLFAINFYDIHAASGNVIKKCRDVMGAGYCGIPGMCGVYKGACAKTCGGCGGRKRIV
GENEEGDMNDQNYNMLNYWRRRDLLRQKDFLQQLRENDEKTLYKM

>TCONS_00069946-protein
MGLVNLLLHLSIIFLVLSATPPLKCHYCHDYEEECLKTNDVTSCPGSQYSCLLHYEETKILNEDGRYDFSRT
VTQDCVVENRNYCENRMRSDPGILSCKAYWCYKDLCNFPASKEEKTQEEITSNISDEESKSEKENSSKGI
IVDKNKKFGQFTKSKAEMLMATKLVDCLTVIIVYKLLVVLLFR

>TCONS_00073533-protein
MKISWQLVSFFWFEFHKTACLEPYKRLSCWTCKKGTGDQCTERVNTVCEGRENVCVTSRITEQTKNWK
GEMLIKERIEKKCHFIAPYQECECIVGGLLGETCELLDCCGQDLCNWRNSKQEKARLCSSSSSINVKPFC
GQFVYVVLIGILMLFSQFVLY

>TCONS_00073545-protein
MTLLRIGSINIFFVVFCWMLITWVSTAPGINPLERFCLDSPEKCPISIDEEALKIFCQINTIKCLQAMRDSIEDL
SHDVQNKEIENDVDSIDDFYSRGLTQKDKHKVNIWDVLGDASEKDYPVKYVEVLTKLAAKQNQQRHKRF
MTLSKLDDGQTQRDYPTKYTDLEIQLAKQKQKRRDEMTTPLQQAMFTKWLLNGNIPATSRKIPFHNAHIL
PRFTSRKDNKWDKANDRVPRKVSPTHSSLTRPSFGGNLNRGNKDYKTNKAVFAEAIDRASGKVPLNHN
SINTLPTFGNKDKTSKAHNDEDEDTAAKNSDKDTMNIMKLSDDNAKNSNSEEEINDSAKVTNDSSNKATKI
WLYKIPSDENSIGVASDHPINFFELKERLAGLPKQVDMKSRNIKKKDDVESLKNDEENTLDEKTIERREQA
MDRIK

**Neuropeptide Precursor Candidates - *Nematostella vectensis***

>XP_001638801.3
MAPRPGTLLLLGILIQVLICTAKSTYKKEIADLLDDNKDTPQFWKGRFSDPQFWKGRFSDPQFWKGRFSD
PQFWKGRFSDPQFWKGRFSDPQFWKGRFSDPQFWKGRFSDPQFWKGRFADELLNGGHHKHHHEEG
EWKRTAGPGRFGREDQGRFGREDQGRFGREDQGRFGREDQGRFGREDQGRFGREDQGRFGREDQ
GRFGREDQGRFGREDQGRFGREDQGRFGREDQGRFGREDQGRFGREDQGRFGREDQGRFGREDQ
GRFGREDQGRFGREDQGRFGREDQGRFGREDQGRFGREDQGRFGREDQGRFGREDQGRFGREDQ
GRFGREDQGRFGRNKIARVIDLDQGRFGRTMDTATKKDTVASMPEQDANPQTRFDGKKRQAAKERSIE
KKSTISSDAKASDAKQS

>XP_032219419.2
MAPRPGTLLLLGILIQVLICTAKSTYKKEIADLLDDNKDTPQFWKGRFSDPQFWKGRFSDPQFWKGRFSD
PQFWKGRFSDPQFWKGRFSDPQFWKGRFSDPQFWKGRFADELLNGGHHKHHHEEGEWKRTAGPGR
FGREDQGRFGREDQGRFGREDQGRFGREDQGRFGREDQGRFGREDQGRFGREDQGRFGREDQGR
FGREDQGRFGREDQGRFGREDQGRFGREDQGRFGREDQGRFGREDQGRFGREDQGRFGREDQGR
FGREDQGRFGREDQGRFGREDQGRFGREDQGRFGREDQGRFGREDQGRFGREDQGRFGREDQGR
FGRNKIARVIDLDQGRFGRTMDTATKKDTVASMPEQDANPQTRFDGKKRQAAKERSIEKKSTISSDAKAS
DAKQS

>XP_048577319.1

MAPRPGTLLLLGILIQVLICTAKSTYKKEIADLLDDNKDTPQFWKGRFSDPQFWKGRFSDPQFWKGRFSD
PQFWKGRFSDPQFWKGRFSDPQFWKGRFSDPQFWKGRFADELLNGGHHKHHHEEGEWKRTAGPGR
FGREDQGRFGREDQGRFGREDQGRFGREDQGRFGREDQGRFGREDQGRFGREDQGRFGREDQGR
FGREDQGRFGREDQGRFGREDQGRFGREDQGRFGREDQGRFGREDQGRFGREDQGRFGREDQGR
FGREDQGRFGREDQGRFGREDQGRFGREDQGRFGREDQGRFGREDQGRFGREDQGRFGREDQGR
FGRNKIARVIDLDQGRFGRTMDTATKKDTVASMPEQDANPQTRFDGKKRQAAKERSIEKKSTISSDAKAS
DAKQS

>XP_048577320.1
MAPRPGTLLLLGILIQVLICTAKSTYKKEIADLLDDNKDTPQFWKGRFSDPQFWKGRFSDPQFWKGRFSD
PQFWKGRFSDPQFWKGRFSDPQFWKGRFSDPQFWKGRFSDPQFWKGRFADELLNGGHHKHHHEEG
EWKRTAGPGRFGREDQGRFGREDQGRFGREDQGRFGREDQGRFGREDQGRFGREDQGRFGREDQ
GRFGREDQGRFGREDQGRFGREDQGRFGREDQGRFGREDQGRFGREDQGRFGREDQGRFGREDQ
GRFGREDQGRFGREDQGRFGREDQGRFGREDQGRFGREDQGRFGREDQGRFGREDQGRFGRNKIA
RVIDLDQGRFGRTMDTATKKDTVASMPEQDANPQTRFDGKKRQAAKERSIEKKSTISSDAKASDAKQS

>XP_048578696.1
MYSITYYIHSQTYKPIEAMESKRLLVILVVCFALVAFAPSTEAQGLRWGREYDESPEKLVPMMKEYIRRQE
ELKKKREFSMN

>XP_032227076.2
MRLYLFVPVFALVLAVEGASDEKRDSKQPPIDLSPAAYFHIRGKRTHNAPPLDLSGPAYFHIRGKRTAKQP
PYLDLGEPSFFHIRGKRTEGPPYIDLTEPSFFHIRGKRSSEQPPLDLGPAYFHIRGKRTKNPPIDLGPAYFH
IRGKRLSGEQPPYLDLTPAYFHIRGKRTQQPPMIDLSEPAFFHIRGRRAVEQPPYLDLTPSYFHIRGKRTE
YPPFLELGQPSYFHIRGRRAEKTTKD

>XP_048576188.1
MRLYLFVPVFALVLAVEGASDEKRDSKQPPIDLSPAAYFHIRGKRTHNAPPLDLSGPAYFHIRGKRTAKQP
PYLDLGEPSFFHIRGKRTEGPPYIDLTEPSFFHIRGKRSSEQPPLDLGPAYFHIRGKRTKNPPIDLGPAYFH
IRGKRLSGEQPPYLDLTPAYFHIRGKRTQQPPMIDLSEPAFFHIRGRRAVEQPPYLDLTPSYFHIRGKRTE
YPPFLELGQPSYFHIRGRRVSRWGSLRIPHQGEKSGENNERLTLLNDASTHARNYCGLWTCLRAAQAR

>XP_048576189.1
MRLYLFVPVFALVLAVEGASDEKRDSKQPPIDLSPAAYFHIRGKRTHNAPPLDLSGPAYFHIRGKRTAKQP
PYLDLGEPSFFHIRGKRTEGPPYIDLTEPSFFHIRGKRSSEQPPLDLGPAYFHIRGKRTKNPPIDLGPAYFH
IRGKRLSGEQPPYLDLTPAYFHIRGKRTQQPPMIDLSEPAFFHIRGRRAVEQPPYLDLTPSYFHIRGKRTE
YPPFLELGQPSYFHIRGRRVSRWGSLRIPHQGEKSK

>XP_032221828.2
MMAIRVLFIVASLLALSSALPWSYFQKRAENTCDTPSFHGKLQMNIRAGDKFIDGRATIDYDHLGKRLKLS
ATLHFDGGSESSFIAFYNYQEGVVVIANPGIGGSGVCHKKDMADIPALIPVPVLYGGSLLKSGSLGLSSDD
LHTALYTGLVGFKTRYFTTVAMETCTPLTSVVYHTSEKDGLEVTTTNYYGIVLEEREPEIFKLPEYCTDTRK
RRQMPNMVPFHGKRQMPNMVPFHGKRQMPNMVPFHGKRQMPNMVPFHGKRQMPNQVPYLG

>XP_032221830.2
MMAIRVLFIVASLLALSSALPWSYFQKRAENTCDTPSFHGKLQMNIRAGDKFIDGRATIDYDHLGKRLKLS
ATLHFDGGSESSFIAFYNYQEGVVVIANPGIGGSGVCHKKDMADIPALIPVPVLYGGSLLKSGSLGLSSDD

LHTALYTGLVGFKTRYFTTVAMETCTPLTSVVYHTSEKDGLEVTTTNYYGIVLEEREPEIFKLPEYCTDTRK
RRQMPNMVPFHGKRQMPNMVPFHGKRQMPNMVPFHGKRQMPNQVPYLG

>XP_048586481.1
MMAIRVLFIVASLLALSSALPWSYFQKRAENTCDTPSFHGKLQMNIRAGDKFIDGRATIDYDHLGKRLKLS
ATLHFDGGSESSFIAFYNYQEGVVVIANPGIGGSGVCHKKDMADIPALIPVPVLYGGSLLKSGSLGLSSDD
LHTALYTGLVGFKTRYFTTVAMETCTPLTSVVYHTSEKDGLEVTTTNYYGIVLEEREPEIFKLPEYCTDTRK
RRQMPNMVPFHGKRQMPNMVPFHGKRQMPNMVPFHGKRQMPNMVPFHGKRQMPNQVPYLG

>XP_032233209.2
MRGVFIVFFGFLLTVVSAKSAEERPTKDVRREIPPQGFRFNQWGKKEIPPQGFRFNQWGKKEIPPQGLRF
NQWGKKEIPPQGFRFNQWGKKEIPPQGFRFNQWGKKEIPPQGFRFNQWGKKEIPPQGLRFNQWGKKEI
PPQGLRFSQWGKKEIPPQGFRFNQWGKKEIPPQGLRFNQWGKKEIPPQGLRFNQWGKKEIPPQGLRFS
QWGKREIPPQGLRFNQWGKKEIPPQGLRFNQWGKKEIPPQGLRFSQWGKKEIPPQGLRFNQWGKKEIP
PQGLRFNQWGKRKLINQVMI

>XP_048577066.1
MRGVFIVFFGFLLTVVSAKSAEERPTKDVRREIPPQGFRFNQWGKKEIPPQGFRFNQWGKKEIPPQGLRF
NQWGKKEIPPQGFRFNQWGKKEIPPQGFRFNQWGKKEIPPQGFRFNQWGKKEIPPQGLRFNQWGKKEI
PPQGLRFSQWGKKEIPPQGFRFNQWGKKEIPPQGLRFNQWGKKEIPPQGLRFNQWGKKEIPPQGLRFS
QWGKREIPPQGLRFNQWGKKEIPPQGLRFNQWGKKEIPPQGLRFSQWGKKEIPPQGLRFNQWGKKEIP
PQGLRFNQWGKRKLINQVMI

>XP_048577067.1
MRGVFIVFFGFLLTVVSAKSAEERPTKDVRREIPPQGFRFNQWGKKEIPPQGFRFNQWGKKEIPPQGLRF
NQWGKKEIPPQGFRFNQWGKKEIPPQGFRFNQWGKKEIPPQGFRFNQWGKKEIPPQGLRFNQWGKKEI
PPQGLRFSQWGKKEIPPQGFRFNQWGKKEIPPQGLRFNQWGKKEIPPQGLRFNQWGKKEIPPQGLRFS
QWGKREIPPQGLRFNQWGKKEIPPQGLRFNQWGKKEIPPQGLRFSQWGKKEIPPQGLRFNQWGKKEIP
PQGLRFNQWGKRKLINQVMI

>XP_048581391.1
MKITPVMSCLVLASALLVTAECYRITDPGDLEKPQENDEPGPPMIKIPVRHGKREFNDDLEDTTLLHYPFR
VARKRELFNAGGPPPIYLPVRPGKRGYYRDNIQETRQYLDPRPGKRSRNH

>XP_048580630.1
MKVLCVTLVLMALVCSVQSRNLRHKDYERELQDYRRDFARALSMVEQDAFLPKPRPGRREQDSSNYEF
PPGFHRPGKKRSFKEEK

>XP_048580631.1
MKVLCVTLVLMALVCSVQSRNLRHKDYERELQDYRRDFARALSMVEQDAFLPKPRPGRREQDSSNYEF
PPGFHRPGKKRSFKEEK

>XP_048580632.1
MKVLCVTLVLMALVCSVQSRNLRHKDYERELQDYRRDFARALSMVEQDAFLPKPRPGRREQDSSNYEF
PPGFHRPGKKRSFKEEK

>XP_032233649.1
MYSCMGISLLLILCFKGSYGEESIDLEPGVDKVPEKVEHGEEGSAKYNINTVTLTGNKATGNKVTNDDKRP
PWPPRPGKRTFRPQESADTPSIFRPGRSVHEDQLLFRPGRANLHKRQGLMFRPGRREDVPKPQIFRPG

RREDIPSDDDQLMFRPGRNEEQLFRPGRRSDVGDEQLLFRPGRRSDLEEDQLIFRPGRRSDVAEEQLFR
PGRRSDVPEAFLDQWTSVRPGRGGYRMPWTYTGNSVNSHVSQKSHTFRQKAVEQEKKKREV

>XP_048579807.1
MALFGHTLVAVLFCLALCHAETKRKAADTTDTENELDASPNVNNDDDNDIKRMPTDTKRQAGAPGLWGK
RDAGPPGLWGKRDAGPPGLCRKRSPKPPGLWGKRQAGAPGLWGKRSAGPPGLWGKRDAGPPGLWG
KRVAGPPGLWGKRQAGAPGLWGREAGAPGLWGKRQAGAPGLWGKREAGAPGLWGREANAPGLWG
KRRAGAPGLWGKREANAPGLWGKRQAGPPGLWGKREANAPGLWGKRQAGPPGLWGKREANAPGLW
GKRQAGPPGLWGKRDEDEDEDMDETNGDPLWGRSADAGPPGLWGRKKRAASPQRDLYGIGLWGRNA
ALMTAEELDLSFKNEEQS

>XP_048579808.1
MALFGHTLVAVLFCLALCHAETKRKAADTTDTENELDASPNVNNDDDNDIKRMPTDTKRQAGAPGLWGK
RDAGPPGLWGKRDAGPPGLCRKRSPKPPGLWGKRQAGAPGLWGKRSAGPPGLWGKRDAGPPGLWG
KRVAGPPGLWGKRQAGAPGLWGREAGAPGLWGKRQAGAPGLWGKREAGAPGLWGREANAPGLWG
KRRAGAPGLWGKREANAPGLWGKRQAGPPGLWGKREANAPGLWGKRQAGPPGLWGKREANAPGLW
GKRQAGPPGLWGKRDEDEDEDMDETNGDPLWGRSADAGPPGLWGRKKRAASPQRDLYGIGLWGRNA
ALMTAEELDLSFKNEEQS

>XP_048579809.1
MALFGHTLVAVLFCLALCHAETKRKAADTTDTENELDASPNVNNDDDNDIKRMPTDTKRQAGAPGLWGK
RDAGPPGLWGKRDAGPPGLCRKRSPKPPGLWGKRQAGAPGLWGKRSAGPPGLWGKRDAGPPGLWG
KRVAGPPGLWGKRQAGAPGLWGREAGAPGLWGKRQAGAPGLWGKREAGAPGLWGREANAPGLWG
KRRAGAPGLWGKREANAPGLWGKRQAGPPGLWGKREANAPGLWGKRQAGPPGLWGKREANAPGLW
GKRQAGPPGLWGKRDEDEDEDMDETNGDPLWGRSADAGPPGLWGRKKRAASPQRDLYGIGLWGRNA
ALMTAEELDLSFKNEEQS

>XP_032226775.1
MTAKLCLPLCIYVFCILHCVSTSSLRVKADNNPSDDDALSRFFRRDCHVGLWCGKKREPAIEQSSRFKSA
DDCPVGLWCGKKRSASKGESIEISHSNKQREQGEEESTVLQKLTDGNTGPPDEVARRMFRRASTSCPP
GLWCGKKRSIPEQNPSGLELQLPYLAGTLPSYEVNDEDAPQSDLKKMFRRASTSCPPGLWCGRKRTVY
SNIGIPPRQSSLKSKFERSCPPGLWCGK

>XP_032220490.1
MPRKSHYLIALDARHYPRLYLEMSNYFLHWVTALAFILCSGLANGKIPKVPIKEVENAPPSTGEGEAGFIP
WFSGKREFNPKNALKNDNAPPGNEEGEAGFIPWFSGKRELIPKIVLQNDNAPPGNGEGEAGFIPWFSGK
REFEPKNALQNDNAPPGNGEGEAGFIPWFSGKRDTTKAAVQNDLVPPSKGAGEQGFIPWFSGKRTVNR
LKEINTENDADANKAPPSGHNEAGFIPWFAGKRKV

>XP_048582935.1
MPRKSHYLIALDARHYPRLYLEMSNYFLHWVTALAFILCSGLANGKIPKVPIKEVENAPPSTGEGEAGFIP
WFSGKREFNPKNALKNDNAPPGNEEGEAGFIPWFSGKRELIPKIVLQNDNAPPGNGEGEAGFIPWFSGK
REFEPKNALQNDNAPPGNGEGEAGFIPWFSGKREFEPKNALQNDNAPPGNGEGEAGFIPWFSGKRDTT
KAAVQNDLVPPSKGAGEQGFIPWFSGKRTVNRLKEINTENDADANKAPPSGHNEAGFIPWFAGKRKV

>XP_048586561.1
MKRRIHAILPVVALCVALSLANIDNGEMVERSGPAQPPLSHLSGKRASTWLRADPEQSIEMSRPLQPVDA
LEKGNSGKILQLDKVPMKMMIRQFGATPPSGIGRKRTNFRRYPPLNAVGKKREFCETRKDIRNNAILD

>XP_032235502.1
MESFVLLAVLTYSLVSCTLVTPQSSKEEVATRWSKDKYDYLPLIPHGESSDESDAFLENILRNDKTTYREK
KGRVVHKGKNPPQPPFRIPIGRKRSNRVVIHTGANPTMAPFRIPGGRKRSNRVVIHTGANPTMAPFRIPIG
RKRSNRVVIHTGANPTMAPLRIPFGRKRSNGVVVHTNPDLSDLDGDTPNVNIERAQKEEC

>XP_032226344.1
MVKFEIFMGLALLLFVSADCLENLLQIKRDSSPPQWKRESTPPQWKRDSSPPQWKRDSSPPQWKRDSS
PPQWKLDSSPPQWKRDSSPPQWKRDSSPPQWKRDSSPPQWKRDSSPPQWKQDLSPPQWKRESTPP
QWKLESTPPQWKRDSSPPQWKRGIFH

>XP_001634396.2
MLSLVALVTLALISPELASGSPFSQMRRSLLEEQCLPGVWGCKRGAPAALNEQRRTASEESDEQCLPGV
WGCKKRTVIQERQCLPGVWGCKRGFDIPIQAKKKQKTPEECPPGLWGCKRGFLGGYYPETSPDYYQQG
MPSEYSEYQQGYPSDYQNLASYYKRRLAIRMRRSVEKEDEQCLPGVWGCKKDKIEAKKSEDQCLPGVW
GCKRDKIAKKQAEEQCPPGLWGCKKDNIEIEKTEETKKSEDQCLPGVWGCKKRSVTSN

>XP_032239630.1
MLSLVALVTLALISPELASGSPFSQMRRSLLEEQCLPGVWGCKRGAPAALNEQRRTASEESDEQCLPGV
WGCKKRTVIQERQCLPGVWGCKRGFDIPIQAKKKQKTPEECPPGLWGCKRGFLGGYYPETSPDYYQQG
MPSEYSEYQQGYPSDYQNLASYYKRRLAIRMRRSVEKEDEQCLPGVWGCKKDKIEAKKSEDQCLPGVW
GCKRDKIAKKQAEEQCPPGLWGCKKDNIEIEKTEETKKSEDQCLPGVWGCKKRSVTSN

>XP_048583863.1
MLSLVALVTLALISPELASGSPFSQMRRSLLEEQCLPGVWGCKRGAPAALNEQRRTASEESDEQCLPGV
WGCKKRTVIQERQCLPGVWGCKRGFDIPIQAKKKQKTPEECPPGLWGCKRGFLGGYYPETSPDYYQQG
MPSEYSEYQQGYPSDYQNLASYYKRRLAIRMRRSVEKEDEQCLPGVWGCKKDKIEAKKSEDQCLPGVW
GCKRDKIAKKQAEEQCPPGLWGCKKDNIEIEKTEETKKSEDQCLPGVWGCKKRSVTSN

>XP_048583864.1
MLSLVALVTLALISPELASGSPFSQMRRSLLEEQCLPGVWGCKRGAPAALNEQRRTASEESDEQCLPGV
WGCKKRTVIQERQCLPGVWGCKRGFDIPIQAKKKQKTPEECPPGLWGCKRGFLGGYYPETSPDYYQQG
MPSEYSEYQQGYPSDYQNLASYYKRRLAIRMRRSVEKEDEQCLPGVWGCKKDKIEAKKSEDQCLPGVW
GCKRDKIAKKQAEEQCPPGLWGCKKDNIEIEKTEETKKSEDQCLPGVWGCKKRSVTSN

>XP_032228536.2
MLSVKLCIVLCVFVIVFEESRAVPQDTSLASSDKLSRKNAGAPCVGKDCEERRGIRAKWYRRCPKGALCS
RTTRKEKSEHRRQNKKSFQSRENDEAPSKFAACKPDLMCKKKSDLPRRIAIRLGYRDAQDIPECPDAFW
CKRRQKSQRRSFSCPPGLWCKRLAEDSTVCHEGFNCKRSVSAERLRAAGYSDSAKGCPPGLWCKRAV
YAFKLNCPPGLWCKRESEDDQRPDRRENGIEQKSVKKGIEVDQRPDRQDSGVYQRLEGDENKPSSDCP
PGLWCKRNNEVPRRSDKRVDRISDCPPGLWCKRGLTRNLAAVLGFKRSDSIANCPPGLWCKRSEKSLD
VDCPLGYKCKRQISDDQGDLTAEYPGYETSRTMPECPRGLVCKRLHNNKNSNNQKKSASLPCPPGLWC

>XP_048584269.1
MYSPSILLLVAMATYSVTLLEALPTHDESSCPPGFWCKRKRSADDGNCPPGFWCKRSSQEQDESWGCP
PGFWCRKKRNVIESEVPEDACPPGFWCKKKRDEIPCPSCREVKDDGCPPGFWCKKKRSFDSGSCPPG
FWCRKRREARAATDLETYSCPPGFWCHRSAMVREDETSAGGCPPGFWCRKRTAPEEQTPNEDTFNW
GCPPGFWCRKRRSVDSDETCPPGFWCKRGVYNDNDDGNCPPGFWCKKKRNIKEDPTSQADCPPGFW
CRKKRSL

>XP_048584270.1
MYSPSILLLVAMATYSVTLLEALPTHDESSCPPGFWCKRKRSADDGNCPPGFWCKRSSQEQDESWGCP
PGFWCRKKRNVIESEVPEDACPPGFWCKKKRDEIPCPSCREVKDDGCPPGFWCKKKRSFDSGSCPPG
FWCRKRREARAATDLETYSCPPGFWCHRSAMVREDETSAGGCPPGFWCRKRTAPEEQTPNEDTFNW
GCPPGFWCRKRRSVDSDETCPPGFWCKRGVYNDNDDGNCPPGFWCKKKRNIKEDPTSQADCPPGFW
CRKKRSL

>XP_048584271.1
MYSPSILLLVAMATYSVTLLEALPTHDESSCPPGFWCKRKRSADDGNCPPGFWCKRSSQEQDESWGCP
PGFWCRKKRNVIESEVPEDACPPGFWCKKKRDEIPCPSCREVKDDGCPPGFWCKKKRSFDSGSCPPG
FWCRKRREARAATDLETYSCPPGFWCHRSAMVREDETSAGGCPPGFWCRKRTAPEEQTPNEDTFNW
GCPPGFWCRKRRSVDSDETCPPGFWCKRGVYNDNDDGNCPPGFWCKKKRNIKEDPTSQADCPPGFW
CRKKRSL

>XP_032223176.1
MKPYQKVLAGLLVVALAVQVVLSQETDGESLKKAEEEFVRRCAVYGCGLFDDFEDKRGLFTKLRLGGSK
GGNRHYFKNRKYFRNNVVKRDANLNKLSVKDLDNE

>XP_032218860.1
MPENLPFHLLIQVQLLLIAAKAPLAAEQCNITTSCRWSNDDLCSVDTNAEDCVLDNNCCWGYQEGNAGK
CFNRTFMEGRIEDCLITPPISCKSQRPWTGDIGSWGDALKGKILEEQFVVSRLQCHDFCTRNARCAAVNV
GPVLMGKRVCELLEATDSVQTQQRAGYSGSILIHEEFRKTYLESGCSTQ

>XP_032218861.1
MPENLPFHLLIQVQLLLIAAKAPLAAEQCNITTSCRWSNDDLCSVDTNAEDCVLDNNCCWGYQEGNAGK
CFNRTFMEGRIEDCLITPPISCKSQRPWTGDIGSWGDALKGKILEEQFVVSRLQCHDFCTRNARCAAVNV
GPVLMGKRVCELLEATDSVQTQQRAGYSGSILIHEEFRKTYLESGCSTQ

>XP_032228088.1
MNKISALLFLGCVVAITWAFPREETMASEGPENETRIEDAEELESKSSYLSKREKLEEITGFAKMKKPYSS
RREKLEEMRAFDRKRQTKEQVICEGPAKLALESYHIDLSKSHLCGENSKRWANDFKSFLRKRELEYHMS
SNYRRNMNACCSPDCDWLLQYLEVDSQVIKCVDKK

>XP_032228089.1
MNKISALLFLGCVVAITWAFPREETSIEDAEELESKSSYLSKREKLEEITGFAKMKKPYSSRREKLEEMRAF
DRKRQTKEQVICEGPAKLALESYHIDLSKSHLCGENSKRWANDFKSFLRKRELEYHMSSNYRRNMNACC
SPDCDWLLQYLEVDSQVIKCVDKK

>XP_032228091.1
MNKISALLFLGCVVAITWAFPREETMASEGPENETRIEDAEELESKSSYSSRREKLEEMRAFDRKRQTKE
QVICEGPAKLALESYHIDLSKSHLCGENSKRWANDFKSFLRKRELEYHMSSNYRRNMNACCSPDCDWLL
QYLEVDSQVIKCVDKK

>XP_032228092.1
MNKISALLFLGCVVAITWAFPREETTYLSKREKLEEITGFAKMKKPYSSRREKLEEMRAFDRKRQTKEQVI
CEGPAKLALESYHIDLSKSHLCGENSKRWANDFKSFLRKRELEYHMSSNYRRNMNACCSPDCDWLLQY
LEVDSQVIKCVDKK

>XP_032228093.1

MNKISALLFLGCVVAITWAFPREETSIEDAEELESKSSYSSRREKLEEMRAFDRKRQTKEQVICEGPAKLA
LESYHIDLSKSHLCGENSKRWANDFKSFLRKRELEYHMSSNYRRNMNACCSPDCDWLLQYLEVDSQVIK
CVDKK

>XP_048578036.1
MHSVVLSMAVFAFLQLNAMFVQGEQSCPAAVKVGCYEDKVHHRALGKLLFQDRSETGDRFSGVLIDWH
HYDKYLEGLICRCATAAKGKGYTHFGLQFYGECHSAPNSLANYGKHGKSDNCVDHRLVECEDETSDHC
VGLSNANYVYRVTGKMIRVLVIFLLATVVEAVNQTCSEDFYKLGCFRDSRWSRGMSELLINDRQRSSPNQ
LNWNDWNKYLPGLACRCAAKAREKGYTLFGIQFYGECWAGAGACDMYDQQGTSLRCVGQDFRSCDN
SDNQACAGGPNANYVYLLTE

>XP_032224051.1
MHPIKTNCERCARCRYFFPSLAMKTVLFFMFLLFEGGDGVICEVCGWNTTSSTYRQCMASLDLYDCDRI
GSTYDRCFISCNTAGTDNWEFGCTTSVQCTRKARSCKSESCQFECCQGDMCTRNMTDPCRGSKQHAP
AVITISTCVVLGSILSFYGRVL

>XP_032224052.1
MKTVLFFMFLLFEGGDGVICEVCGWNTTSSTYRQCMASLDLYDCDRIGSTYDRCFISCNTAGTDNWEFG
CTTSVQCTRKARSCKSESCQFECCQGDMCTRNMTDPCRGSKQHAPAVITISTCVVLGSILSFYGRVL

>XP_032227339.2
MHLQLSIAVVVVLLATSCVGGKVGEYTTSAGHMKNLLKHAHLESHGFTRKDVMESRQVPGCKEPVDQVK
SVYYKSVQCHKAAEVEKQLNRLMKGVFGKKTNTQVARKTSALGRGEICEVGMEVDFNSKTTCQPETPK
RAEAKGTATSRLIIIIIFG

>XP_001641384.1
MDLPMAVLVSLLVASVVIASPVTKGKHQGTSWMQCKTSRDCEMDECCVALMKSEVKVCKRRPQLGNHC
TPTIMPGIDGRCPCSSGLTCALTFMDNMGLRDKYQCVLVPDSDREFEKRRQ

>XP_032235296.2
MRAFITCIAFFLVVGLVFGEERDESNIKASEKSDKLPKGAPSLPDAVAAKKCKPMFAACRKQSPEKVRECF
TAFYNGCFLKVRNMLNNIEKCLSAADQEIVDAYMAARLEQSMGGE

>XP_048589131.1
MRAFITCIAFFLVVGLVFGEERDESNIKASEKSDKLPKGAPSLPDAVAAKKCKPMFAACRKQSPEKVRECF
TAFYNGCFLKVRNMLNNIEKCLSAADQEIVDAYMAARLEQSMGGE

>XP_032222229.2
MSNLCLRRCFAVSKAAVFSAAIAILLSSVMGTPLVDSKSSLNEIGNFEMVKKDIRIVKGQDQKGFFMEQHV
NRPTHRVLRAPSNLFLSQIRPKFPNHIPPCATTEQTLDPANYFNTPHTFVPDSYNRVECSGRVGARGLPE
CVYGIMDCVTVYKDLHFIRKPSHDCSTWYDDVIPNVPSGCMCMWPN

>XP_032222230.2
MAIYKAAVFSAAIAILLSSVMGTPLVDSKSSLNEIGNFEMVKKDIRIVKGQDQKGFFMEQHVNRPTHRVLR
APSNLFLSQIRPKFPNHIPPCATTEQTLDPANYFNTPHTFVPDSYNRVECSGRVGARGLPECVYGIMDCV
TVYKDLHFIRKPSHDCSTWYDDVIPNVPSGCMCMWPN

>XP_032225251.1

MSIAQSLAKRRTYFNAKMASLQYLSVCTSLLLLVCFLNVGRVSSTEVKSGQLYQLCGDQFLSAYTQCCEN
PAGCEGSTRKREMKEGDKSILSPETEAKRFLQFSIARDRVRRGILSKRSAGGGTNPVEECCSETCSLAEV
AEYDCV

>XP_032225252.1
MASLQYLSVCTSLLLLVCFLNVGRVSSTEVKSGQLYQLCGDQFLSAYTQCCENPAGCEGSTRKREMKEG
DKSILSPETEAKRFLQFSIARDRVRRGILSKRSAGGGTNPVEECCSETCSLAEVAEYDCV

>XP_032225253.1
MASLQYLSVCTSLLLLVCFLNVGRVSSTEVKSGQLYQLCGDQFLSAYTQCCENPAGCEGSTRKREMKEG
DKSILSPETEAKRFLQFSIARDRVRRGILSKRSAGGGTNPVEECCSETCSLAEVAEYDCV

>XP_032225254.1
MASLQYLSVCTSLLLLVCFLNVGRVSSTEVKSGQLYQLCGDQFLSAYTQCCENPAGCEGSTRKREMKEG
DKSILSPETEAKRFLQFSIARDRVRRGILSKRSAGGGTNPVEECCSETCSLAEVAEYDCV

>XP_048579001.1
MNTQVRIVLVVAMTTMLVIACECGDLRSYYCEMNTVDKIEMFFRFNCRQQMHKLWGDEAAKSAKPKLSR
SRLSKRECLTMRDYADFYQDCCLSERGCDPEYYLQSYCNCY

>XP_032223656.1
MWRELILLACLLGLGCAAKKSQHESTDDAVKRSIAMSSKSLSDARDDDIDHYTRDDAYAALAFYDYIMKR
DSQLAKRIECKDKFPVFCRKAVAFCLMPMYKVHGNGRLRRKYNIARKFCPVSCHNNGC
>XP_032220484.2
MATFSLIFLAMFAASSGASLYENTNLPTCSEIRGDSCEDDAQCEQCVPDLLSDRKLICNWKWKCNKMSIID
IVSRQQCKNVFKKDCTIDAECLCRETPMVCENRECVKAKSHKVAARL

>XP_032233759.2
MKLLPVVVVALMVVVMEKWLSVQASSEEVEKELKQLEEQLSLGSDTSPKETEIASDETKRGLRAPCGFSP
ACTRRRRRSSRRSFWGKKRSEVESKAQFSDVGTDDLVHVLEDIEKIRHELYDIEAHENTHVTRDEKHVG
KTRDERDANQGSDRA

>XP_032233732.2
MKNLLLKTVIRLIFAKFATGFQCHDTTNSGIYVCDVGSRGACVTVELAATNISSGAQRTVTQRLCIADHSC
NVTRICDGISFSMQQWNISRCHAKCCNTSLCNAPQPTSTVSSTTTEVEQRSQPEVAGTHVVTAPRPLCY
KCPPGTSAETCTQLPSTVQCESSESVCFSVIGKDRIGEIGYRGCTDTEHCDASKLCNQASRDLFLKTGEG
LQDCNGYCCTGEMCNRFNPTRPYNREQETGSTERDIKVVRRETPASSNPPINCYRCVPDNLGSLCKREA
RASRCPLDRSRTGYDGCFSMTGVITNSSTGMVKDEGIWKDCSVVSADCNENGTRCDRLTSVLKDRGLT
LKNCYVNCCQGDLCNNFVPSPTPEVMMARSGTRQCTVGCTRVLLPVLLVVMVTSGYK

>XP_048583936.1
MLAIFLIVAFFSATAKSSSIPVSSFQRLEATHGGNCLHVFYKVCDPPEACKVEFTAKFEDGSSTVKTSAGK
MKPTITPDNSSYVFCMYDLTSSINASCCKYNLSMIQGYGISRHDYISFQVEMCIVQSNVSSFCGTGGWQK
FVVPSKNPCLLEKPCRNFGTCSPAPNDYNCTCRSDGLVTGKNCDVPKNYIKLSNQCLMSNESIATKINQTI
LECAAACDVDANCNSMEFTRISSDSDIGDCVLRKATRSTSATTQCLSDYFEMKTGEKPTADASRGTASV
QIIFALISLGLGFSV

>XP_032233169.1

MKVEVAVFSALVLVANLAIEAESWMCSPYNCPRRRSKIAGVTREPSRTDYRLTACPDGKGLCLRRFRDIF
RDDDSSEKIDGTSLQTRAQPSATDALTRRIEALRMDKDAMREIQTRLRIKSAADQIGE

>XP_032220009.2
MISRSVFVLGLLSLAVVAFAEQEDASKHSALPKSIKQSKKRTCLSWDCADGDESEGLQVVIAEPPYEEEY
EDDSHERAGSENGDVRYPVLEDIFGKRAKKGGSRRCLGWMCREKRAKEQDKKRKVSPQRRRTVKSGS
GSSIFPVKRSKLTKKCLGWDCHNYRRSDDKKRMEARKGARREEQAKKREGNETQQKGNKRCLGWKCS
YYYKKKRGMARDTGSREWRRTVKEAAQKRSTGLKNDGVIPI

>XP_048584370.1
MTKLKTQSKRKSAAQHSKLTMTSFAVSFPLVFLASSLLSHYPVDSSNVIITTDDGSKDKAYCMYEDMMFQ
CKTEKECDGEDYVDWGRGKGQGNDPNLKQNDCKYPQTCCLPFKNSEDSVERYKKKVAADKGKSDAMF
A

>XP_032223012.1
MSAGKDTSTQKFFTGSSYHTMTRATVFFVLLGIIVQFGHKKAEKQTTKPTCSFGTKIYESGDVFTPNSCQ
DICSCKDDGQFHCVPCQAKQPDTNKGCKGWSKHVVVVDSYIGTKKHGCHCYKYVCKRKTIKTVIKELWE
KLKTKFSSLVHPHKPSRDWG

>XP_001638561.1
MAVSLMLLGFLLLGSDSVLGSSREELDPRPRKGPLVHGLVKEAETGAKVEHWKRDTGSSGIPAWMKDT
GKAYNGYPSQVTCALKPMSEAYHELKARRYQNCDQYFRCQGSKKAAKCGLFSKLLASAISYDYRWRTA
GLETYTSAAD
