## Supplementary Figures for "Identifying neuropeptides in *Hydra*: a custom pipeline reveals a non-amidated regulator of muscle contraction and other new members"

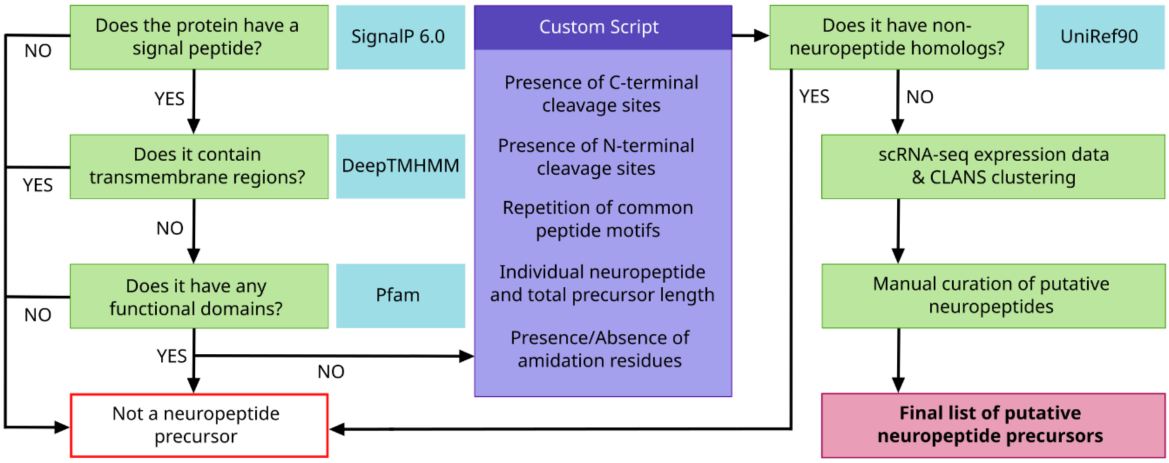


**S1 Figure.** A comprehensive flowchart of the neuropeptide prediction pipeline briefly described in Figure 1A.

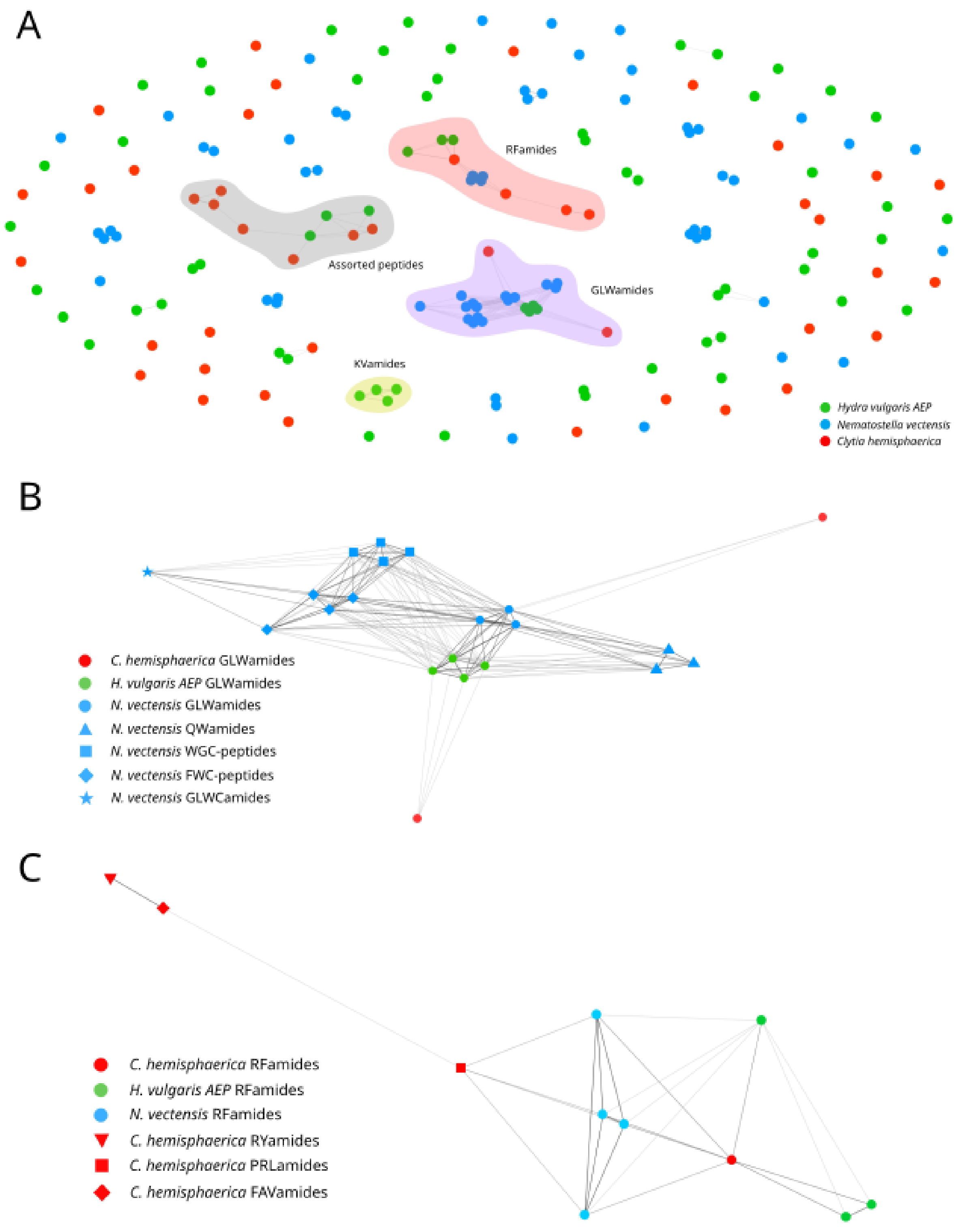


**S2 Figure. CLANS similarity clusters of predicted candidates from three cnidarian species.** (A) Complete CLANS map of all predicted candidates. (B) Cluster containing Wamide sequences and their derivatives. (C) Cluster containing RFamide sequences and their derivatives.

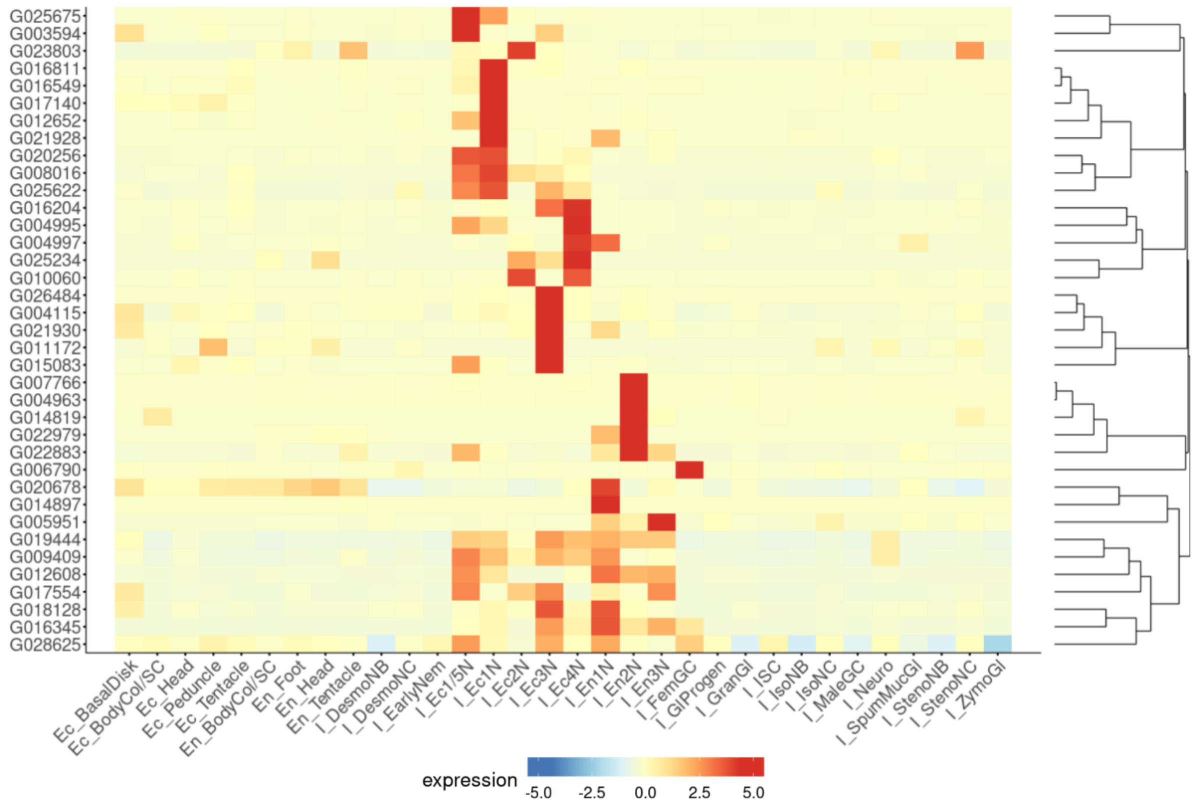


**S3 Figure.** Heatmap of expression of all *Hydra* neuropeptide gene candidates excluding previously reported precursors in single-cell neuronal clusters.

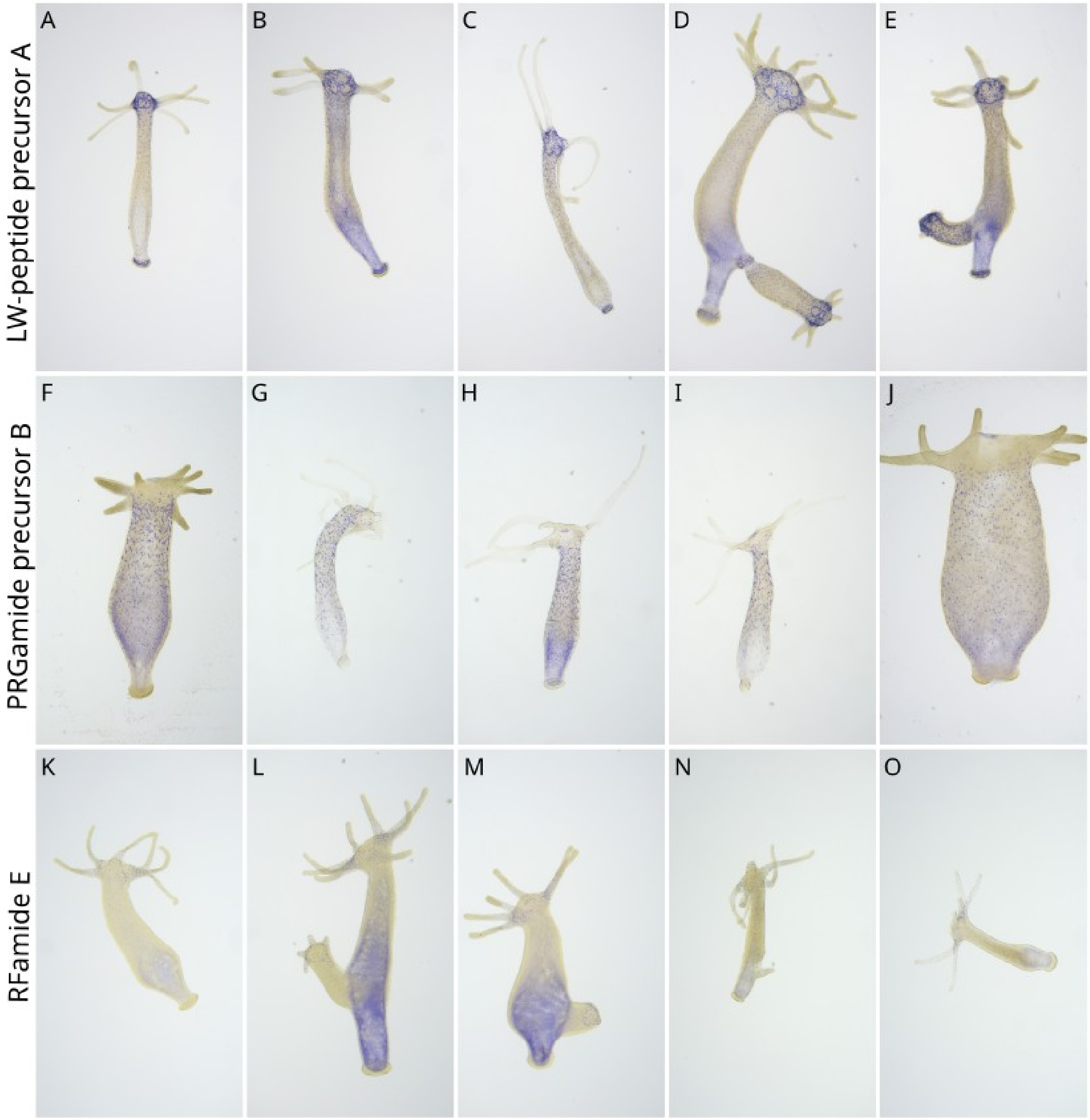


**S4 Figure. Additional whole mount *in situ* hybridization images of the three new *Hydra* neuropeptide genes.** (A-E) LW-peptide precursor A. (F-J) PRGamide precursor B. (K-O) RFamide Preprohormone E.

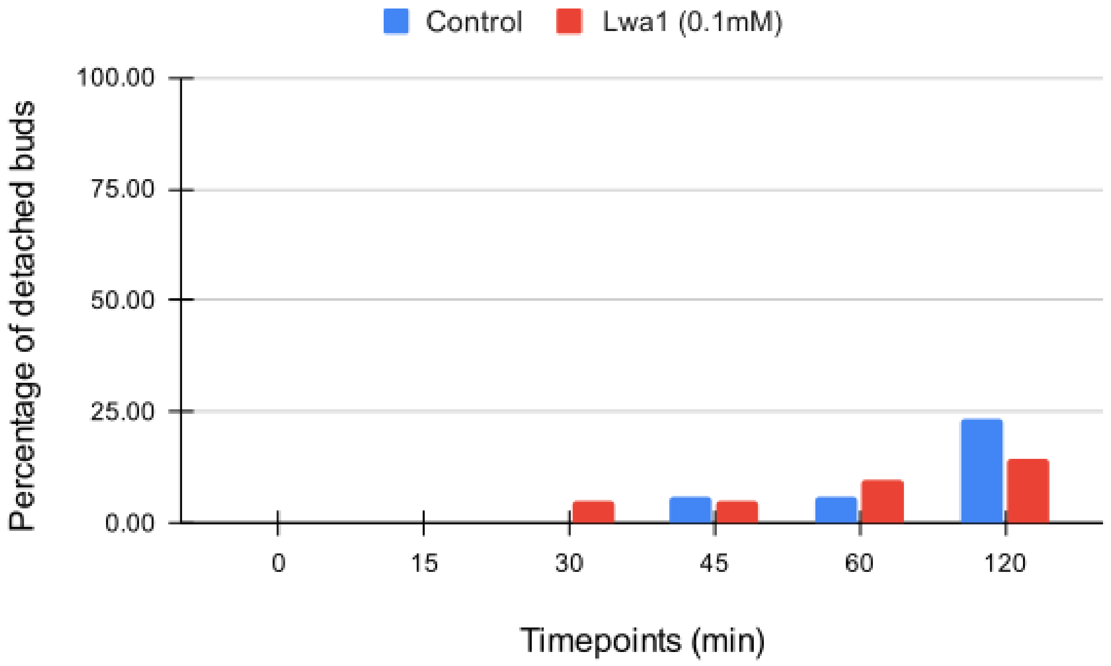


**S5 Figure.** Results of the bud detachment assay between control and LWa1-treated polyps showed no discernible difference in effect.
