## Supplementary material for "Identifying neuropeptides in *Hydra*: a custom pipeline reveals a non-amidated regulator of muscle contraction and other new members": S1_File

#NEUROPEPTIDE PREDICTION SCRIPT

from Bio import SeqIO
import csv
import re

#a function to find a given list of motifs in a sequence
def find_motifs(sequence, motif_list):
    matches = {}
    for motif in motif_list:
        matches[motif] = [match.start() for match in re.finditer(motif, sequence)]
    return matches

#the main function of this script that parses the entire fasta file and generates the motif count and peptide output
def main_function(fasta_file, c_term_motifs, n_term_motifs, aa_1, aa_2, output_file):

#opening the output file to write output   
    with open(output_file, mode = 'w', newline = '') as csv_file:
        fieldnames = ['Sequence ID','Number of C-terminal sites','Putative Peptides']
        writer = csv.DictWriter(csv_file, fieldnames=fieldnames)
        writer.writeheader()

#parsing the fasta file and removing sequences with length >500aa and with no motifs founds
        for record in SeqIO.parse(fasta_file, "fasta"):
            sequence = str(record.seq)
            if len(find_motifs(sequence, c_term_motifs)) == 0 or len(sequence) > 500:
                continue

#find and count the number of c-terminal motifs           
            c_term_matches = find_motifs(sequence, c_term_motifs)
            c_term_motif_count = sum(len(matches) for matches in c_term_matches.values())
            peptides = []

#find n-terminal motifs within a given aa range before each c-terminal motif
            for motif, matches in c_term_matches.items():
                for c_term_match in matches:
                    start_index = max(0, c_term_match - aa_1)
                    end_index = c_term_match - aa_2
                    n_term_matches = find_motifs(sequence[start_index:end_index], n_term_motifs)
                    if any(n_term_matches.values()):

#if n-terminal motif found, extract the encoded putative peptide
#peptide conditions: should not contain multiple repeats of listed motifs within it and should be more than 4aa long
#writing output headers and rows           
                       last_n_term_match = max(n_term_matches.values())[0]
                       peptide = sequence[start_index + last_n_term_match:c_term_match]
                       has_repeats = False
                       for motif in c_term_motifs + n_term_motifs:
                           if peptide.count(motif) >= 3:
                               has_repeats = True
                               break
                       if len(peptide) >= 4 and not has_repeats:
                           peptides.append(peptide)
            if peptides and c_term_motif_count > 0:
                writer.writerow({'Sequence ID': record.id, 'Number of C-terminal sites': c_term_motif_count, 'Putative Peptides': peptides})

#conditions and parameters           
fasta_file = "script_input_aep.fasta"
c_term_motifs = ['KK','KR','RK','RR','GK','GR']
n_term_motifs = ['Q','P','KK','KR','RK','RR']
aa_1 = 20
aa_2 = 2
output_file = "peptides.csv"

main_function(fasta_file, c_term_motifs, n_term_motifs, aa_1, aa_2, output_file)
